# The Ly6/uPAR protein Bouncer is necessary and sufficient for species-specific fertilization

**DOI:** 10.1101/354688

**Authors:** Sarah Herberg, Krista R. Gert, Alexander Schleiffer, Andrea Pauli

## Abstract

Fertilization is fundamental for sexual reproduction, yet its molecular mechanisms are poorly understood. Here, we identify an oocyte-expressed Ly6/uPAR protein, which we call Bouncer, as a crucial fertilization factor in zebrafish. We show that membrane-bound Bouncer mediates sperm-egg binding and is thus essential for sperm entry into the egg. Remarkably, Bouncer is not only required for sperm-egg interaction, but also sufficient to allow cross-species fertilization between zebrafish and medaka, two fish species that diverged over 150 million years ago. Our study thus identifies Bouncer as a key determinant of species-specific fertilization in fish. Bouncer’s closest homolog in tetrapods, SPACA4, is restricted to the male gonad in internally fertilizing vertebrates, suggesting that our findings in fish have relevance to human biology.

## Introduction and Results

Fertilization, whereby two gametes fuse to form the single-cell zygote in sexually reproducing organisms, is highly efficient yet species-restricted. This strategy ensures reproductive success and the survival of distinct species, but how nature has fulfilled these seemingly contradictory requirements is far from understood, particularly at the molecular level. The only vertebrate proteins known so far to be essential for sperm-egg binding are the sperm-expressed IZUMO1 (*1*, *2*) and the egg membrane proteins JUNO (*3*) and CD9 (*4–6*). Binding of IZUMO1 to JUNO mediates adhesion between sperm and egg in mammals (*1–3*, *7*), but the role of CD9 in this process is still unclear. Moreover, none of these factors seem to play a role in mediating species-specificity of fertilization (*8*).

To identify candidate factors required for fertilization in vertebrates, we examined our collection of predicted protein-coding genes (*9*) hat are expressed in zebrafish oocytes and/or testis. A single-exon gene stood out due to its high expression in zebrafish oocytes (Fig. 1A) and the presence of homologous sequences in other vertebrates. We named this gene *bouncer* (*bncr*) based on its loss-of-function phenotype (see below). *Bouncer* lacks any gene annotation in the newest zebrafish genome release (GRCz11). However, our RNA seq and *in situ* hybridization analyses, as well as ribosome profiling data (*9–11*) and CAGE-based transcription start site analysis (*12*), suggested that *bouncer* is a maternal transcript that generates a 125 amino acid (aa) protein (Fig. 1A; Supplementary Fig. 1A, B). *Bouncer* encodes an N-terminal signal peptide and a putative C-terminal transamidase cleavage site, suggesting that the 125 aa protein is processed into an 80 aa glycosylphosphatidylinositol (GPI)-anchored mature protein (Fig. 1A). Consistent with two predicted N-glycosylation sites (Fig. 1A), a Bouncer-specific antibody detected glycosylated Bouncer in the oocyte (Fig. 1B). After deglycosylation, Bouncer is ~10 kDa, as expected for an 80 aa protein.

**Figure 1:**
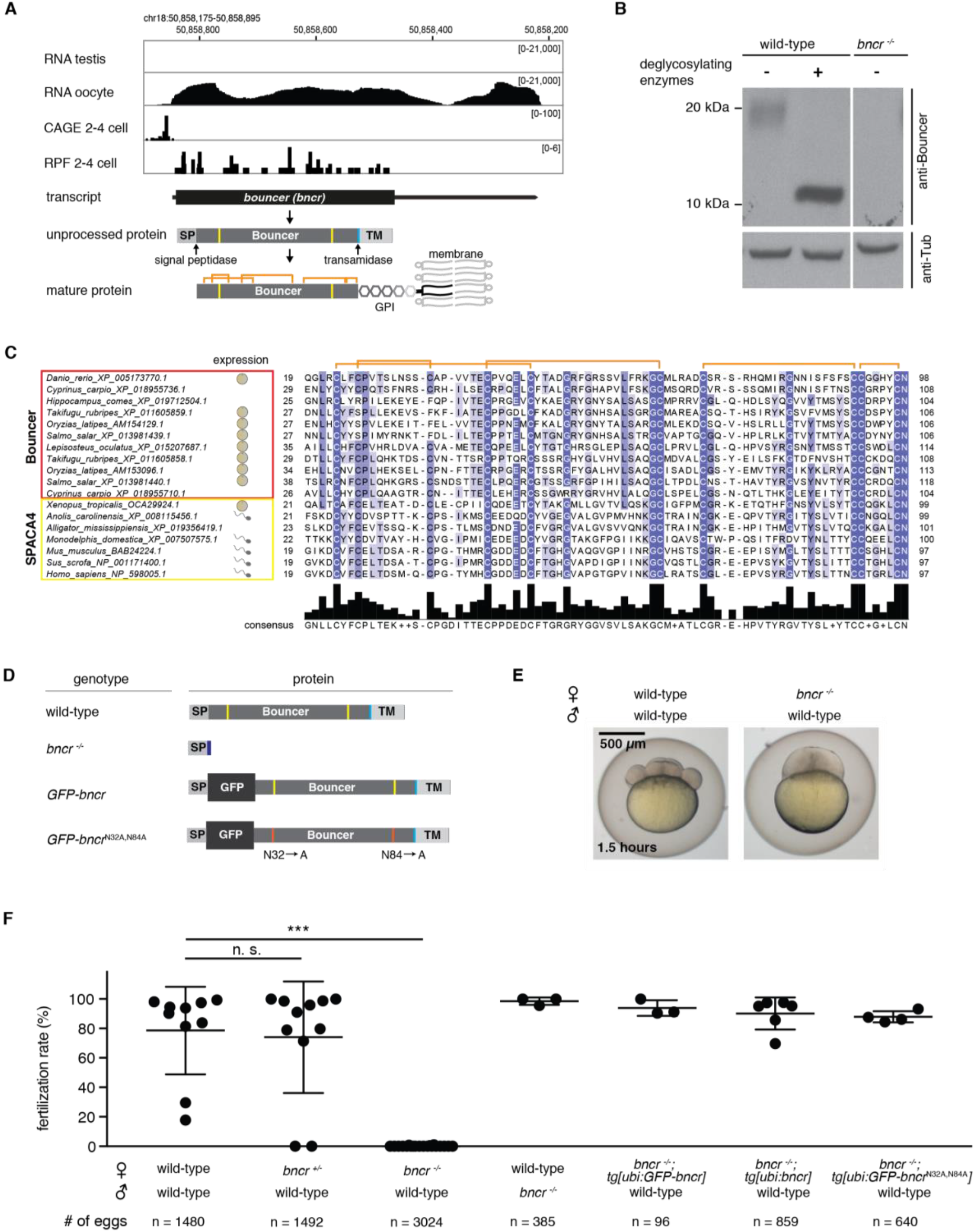
Identification of Bouncer in fish. A – Expression and genomic features of Bouncer. Coverage tracks for RNA sequencing, ribosome profiling (Ribosome Protected Fragments (RPF)) (*10*) and CAGE data (transcription start sites) (*12*) show Bouncer to be expressed in oocytes, but not in testis. Genomic coordinates are based on GRCz10. Bouncer is predicted to encode a short protein containing a signal peptide (SP), an 80-amino acid-long mature domain with ten cysteines and two predicted N-glycosylation sites (yellow), and a transamidase cleavage site (turquoise) followed by a transmembrane region (TM). Mature Bouncer is predicted to be transferred to a GPI-anchor. Orange brackets indicate predicted disulfide bonds. B – Endogenous Bouncer protein is glycosylated. Oocyte-expressed, endogenous Bouncer protein runs at a higher molecular weight (~20 kDa) than predicted by the size of its mature protein (10 kDa), but shifts down to the expected size after treatment with deglycosylating enzymes. Twenty one-cell-stage embryos without yolk were loaded per lane. Bouncer was detected with a Bouncer-specific antibody. No signal is detected in oocytes from *bncr*^−/−^ mutant females, confirming the specificity of the antibody. Tubulin is shown as a loading control. C – Protein sequence alignment of members of the Bouncer/SPACA4 protein family. Apart from the well conserved cysteines (orange brackets indicate predicted disulfide bonds), Bouncer/SPACA4 shows high amino acid divergence among different species. Only amino acid sequences of the conserved mature domain are shown; the extent of the mature domain displayed here is based on the prediction for zebrafish Bouncer. For all species for which expression data was available ((*28–32*); human expression based on GTEx Portal; ESTs based on NCBI), *bouncer/Spaca4* RNA was found to be restricted to either the male (symbol: sperm) or female (symbol: egg) germline. For sequences and accessions see Supplementary Datasets 1 and 2 and Supplementary Table. D-F – Lack of Bouncer causes near-sterility in female zebrafish. *bncr*^−/−^ females lack mature Bouncer protein and are nearly sterile, while *bncr*^−/−^ males are fertile. Transgenically expressed untagged (*tg[ubi:bncr]*), GFP-tagged (*tg[ubi:GFP-bncr]*) and non-glycosylatable Bouncer (*tg[ubi:GFP-Bncr*^*N32A,N84A*^]) rescue the mutant phenotype. (D) Schemes of Bouncer protein products (similar as in (A)); the additional seven amino acids of the *bncr*^−/−^ mutant before the premature stop codon are indicated in dark blue. (E) Representative images show the development of embryos derived from crossings of wild-type and *bncr*^−/−^ females to wild-type males after 1.5 hours, when wild-type embryos have progressed to the eight-cell stage. (F) Quantification of the fertilization rates of the indicated crosses. n = total number of eggs. Mean and standard deviation are indicated (significance tested by Kruskal-Wallis with Dunn’s test for multiple comparisons using statistical hypothesis testing; n. s. > 0.999; *** < 0.0001).

A protein domain search classified Bouncer as a member of the Ly6/uPAR (Ly6/urokinase-type Plasminogen Activator Receptor) protein superfamily, which includes proteins as diverse as toxins, immunoregulators, and cell surface receptors. This protein family is characterized by a 60-80 aa domain (Pfam domain UPAR_LY6_2; E-value 1.4e-05) containing 8-10 highly conserved, distinctly positioned cysteines that form a three-finger type structure (Fig. 1A, C; Supplementary Fig. 2A) (*13*). Apart from the cysteines, the other amino acids have diverged substantially between and within individual subgroups of this large protein superfamily (Fig. 1C; Supplementary Figure 2A, B). BLASTP searches with zebrafish Bouncer and phylogenetic sequence analyses suggested that SPACA4 is the closest homolog in mammals, reptiles, and amphibians (Fig. 1C; Supplementary Figure 2A-C; Supplementary Dataset 1, 2 and Supplementary Table). Human SPACA4/SAMP14 (Sperm Acrosome Membrane-Associated Protein 4/Sperm Acrosomal Membrane Protein 14) was originally identified in a proteomics study as a sperm acrosomal protein, and *in vitro* experiments implied a possible function in fertilization (*14*). However, the *in vivo* function and importance of SPACA4 are unknown.

Intrigued by our finding that Bouncer is expressed in zebrafish oocytes, and that its closest homolog in humans is reported to be expressed in sperm (*14*), we analyzed the expression patterns of other Bouncer/SPACA4 homologs using published expression data. Consistent with the data from zebrafish and humans, we found that externally fertilizing vertebrates (e.g. fish and amphibians) show oocyte-restricted expression of this Ly6/uPAR family member, whereas all internally fertilizing vertebrates we analyzed (e.g. reptiles and mammals) show testis-specific expression (Fig. 1C; Supplementary Figure 2A). Together, these results identified Bouncer and SPACA4 as homologs with opposing sex-specific, germline-restricted expression patterns in internally versus externally fertilizing vertebrates.

To investigate the function of Bouncer, we used CRISPR/Cas9 to generate *bouncer* knockout zebrafish. We established a stable mutant line with a *bouncer* allele that carries a 13-nt deletion, which causes a frameshift at the end of the signal peptide sequence. Eggs from homozygous *bncr*^-/-^ fish lacked mature Bouncer protein, as expected (Fig. 1B, D; Supplementary Fig. 3A). In-crosses of *bouncer* heterozygous (*bncr*^+/−^) fish gave rise to homozygous mutant adults (*bncr*^−/−^) at a Mendelian ratio of ~25%, suggesting that Bouncer is not essential for embryonic or larval development. However, *in vivo* mating experiments showed that only 7 out of 3024 (0.11%) of *bncr*^−/−^ eggs developed into cleavage-stage embryos, as opposed to the majority of eggs from wild-type or *bncr*^+/−^ females (Fig. 1D-F; Supplementary Fig. 3A). In contrast, the reproductive competence of *bncr*^−/−^ males was indistinguishable from wild-type males (Fig. 1F). Notably, female near-sterility was fully rescued by ubiquitous expression of transgenic Bouncer (tg[*ubi:bouncer*]) or GFP-tagged Bouncer (tg[*ubi:sfGFP-bouncer*] or tg[*actb2:sfGFP-bouncer*]) (Fig. 1D, F; Supplementary Fig. 3A, B), confirming that the observed defect was indeed due to the lack of Bouncer protein. Interestingly, ubiquitous expression of a Bouncer mutant that cannot be glycosylated (tg[*ubi/:sfGFP-bncr*^N32A,N84A^]; Supplementary Fig. 3A, C) also fully rescued female near-sterility (Fig. 1F), demonstrating that glycosylation of Bouncer is neither required nor contributes to the essential function of Bouncer. Thus, oocyte-expressed Bouncer is necessary for efficient reproduction in zebrafish.

Zebrafish eggs are activated by the first contact with the spawning medium, independently of the presence of sperm. We considered that eggs from crosses of *bncr*^−/−^ females might remain undeveloped due to defective activation. However, eggs produced by *bncr*^−/−^ females did not show any defects in spawning medium-induced egg activation, as evident by normal chorion (the outer protective shell of fish embryos) elevation, polar body extrusion, and cytoplasmic streaming (Supplementary Fig. 4A-C; Supplementary Movie 1). Moreover, morphological analysis of the chorion revealed that the micropyle, an opening that serves as the sole entry point for sperm into fish oocytes, is present in *bncr*^−/−^ eggs and of similar size as in wild-type eggs (Supplementary Fig. 4D). Taken together, these results suggest that Bouncer is not required for egg activation.

Because eggs from *bncr*^−/−^ females lack any apparent morphological defects, yet do not develop beyond the one-cell stage, we reasoned that Bouncer might be required for fertilization and/or the initiation of early embryonic cleavage cycles (Fig. 2A). To distinguish between these possibilities, we first asked whether sperm can enter eggs that lack Bouncer. To this end, we performed *in vitro* fertilization (IVF) of wild-type (control) and Bouncer-deficient eggs with MitoTracker-labeled sperm and counted the number of eggs with a sperm-derived MitoTracker signal. We detected sperm in about 50% of control (wild-type) eggs (Fig. 2B), consistent with the obtained IVF rates of 50-60% for wild-type eggs. In contrast, we never detected the MitoTracker label in Bouncer-deficient eggs (Fig. 2B), suggesting that Bouncer might play a role in sperm entry during fertilization. If Bouncer’s sole function is to allow sperm to enter the egg, then delivery of sperm into Bouncer-deficient eggs by intra-cytoplasmic sperm injection (ICSI) should restore embryonic development beyond the one-cell stage. Indeed, intra-cytoplasmic injection of wild-type sperm into wild-type or Bouncer-deficient eggs resulted in similar rates of embryos undergoing early embryonic cleavages (Fig. 2C). Thus, delivery of sperm into the egg can bypass the requirement for Bouncer. These results reveal that Bouncer’s key function is in fertilization, and that it enables sperm entry into the egg.

**Figure 2:**
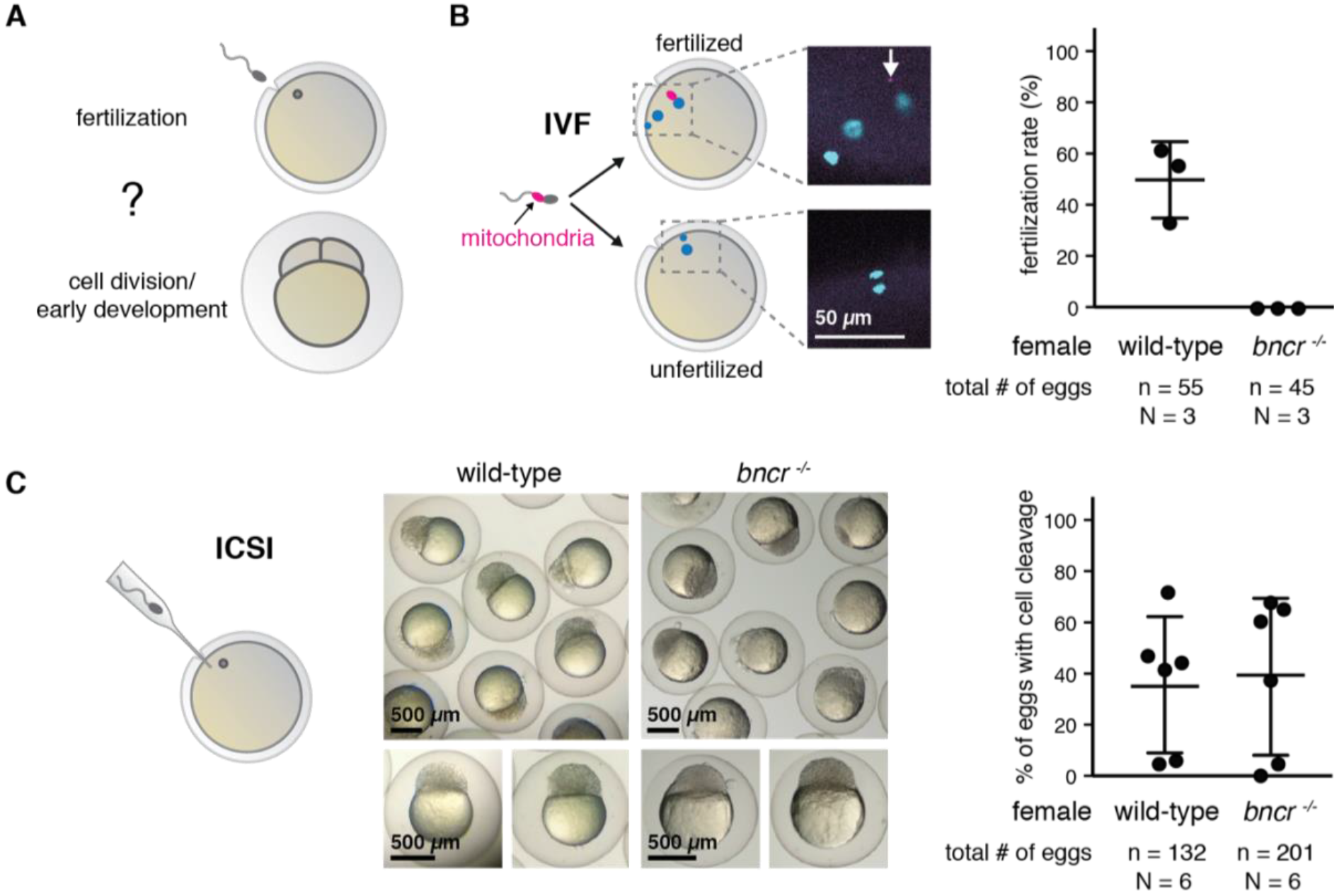
Bouncer is required for sperm entry into the egg. A – Bouncer could either be required for fertilization or for cell division/early development. B – Sperm does not enter *bncr*^−/−^ eggs. Left: Experimental setup. Wild-type sperm was stained with MitoTracker label and used for *in vitro* fertilization (IVF) of wild-type and *bncr*^−/−^ eggs. The eggs were fixed and DAPI-stained before imaging. Representative images are shown; the arrow points to the red MitoTracker signal. Right: Percentage of fertilized eggs, as indicated by the presence of one MitoTracker-labeled sperm and three DAPI signals, deriving from the male nucleus, female nucleus, and the polar body. Mean and standard deviation are indicated (n = total number of eggs; N = number of biological replicates). C – Intra-cytoplasmic sperm injection (ICSI) is able to rescue *bncr*^−/−^ eggs. Left: Experimental setup. Wild-type sperm was injected into wild-type or *bncr*^−/−^ eggs. Injected eggs were scored for cell cleavage after three hours. Representative images are shown. Right: Percentage of eggs that show cell cleavage. Mean and standard deviation are indicated (n = total number of eggs; N = number of biological replicates).

To gain insight into the function of Bouncer during fertilization, we first assessed Bouncer’s localization in oocytes using the fully functional, transgenic GFP-tagged Bouncer (Fig. 1D). Confocal imaging revealed that Bouncer localizes to the egg membrane and to vesicles within the egg (Fig. 3A), consistent with its predicted GPI-anchorage. Compared to the even distribution of a membrane-tethered control fluorescent protein (lyn-Tomato), membrane-localized Bouncer was enriched at the micropyle (Fig. 3A, B). Further, ubiquitous expression of a secreted version of Bouncer lacking the C-terminal membrane anchor (tg[*ubi:sfGFP-bncr*^no™^]) did not rescue the near-sterility of *bncr*^−/−^ females (Supplementary Fig. 3A, B), suggesting that membrane-localization of Bouncer is required for function.

**Figure 3:**
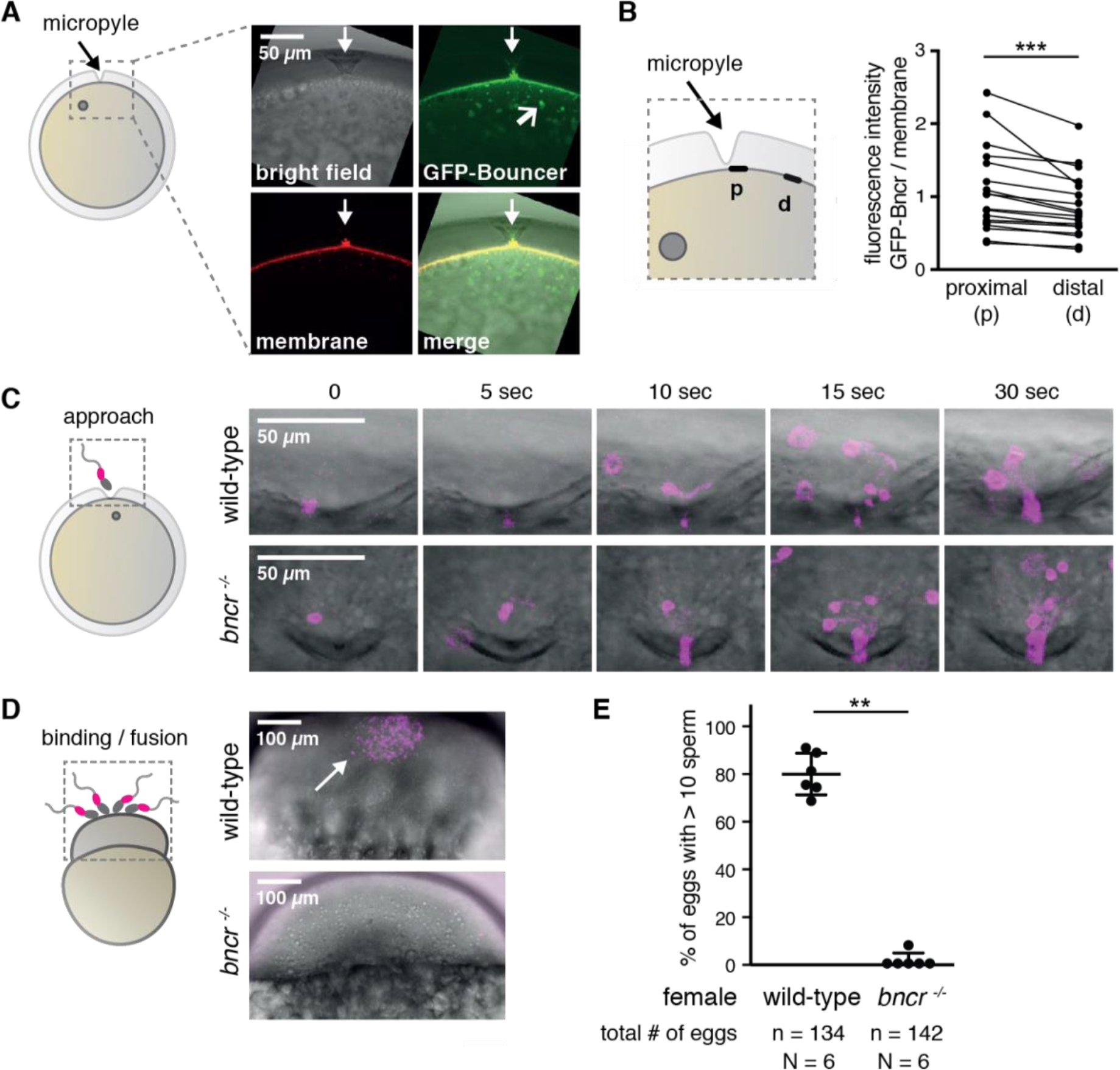
Bouncer mediates binding between sperm and egg. A – Bouncer localizes to the oocyte membrane and to vesicles. Confocal images of eggs expressing GFP-tagged Bouncer (green) and lyn-Tomato (membrane, red) during egg activation show that Bouncer localizes to the egg membrane and to vesicles (open arrow). The position of the micropyle, the single opening in the chorion through which the sperm can enter, is indicated with a filled arrow. Grey circle: oocyte nucleus. B – Bouncer is enriched at the site of sperm entry. The membrane-localized average fluorescence intensity of GFP-Bouncer and lyn-Tomato was measured proximally (p; 1-20 μm) and distally (d; 110-130 μm) from the micropyle for each egg. The paired data points represent measurements from one egg (p = 0.0008, paired t-test; n = 18 (total number of eggs); N = 3 (number of biological replicates)). Grey circle: oocyte nucleus. C – Bouncer is not required for sperm approach. Left: Experimental setup. The approach of MitoTracker-labeled wild-type sperm to the micropyle of unactivated wild-type and *bncr*^−/−^ eggs was monitored by time-lapse imaging. Right: Representative time series of multiple sperm (magenta) approaching the micropyle area of wild-type (top) and *bncr*^−/−^ (bottom) eggs. D – *bncr*^−/−^ eggs are impaired in sperm-egg binding. Left: Experimental setup. Activated and dechorionated wild-type and *bncr*^−/−^ eggs were incubated with MitoTracker-labeled wild-type sperm. Attachment of sperm to the egg membrane was assessed after gentle washing by confocal imaging. Right: Representative images of a wild-type egg (top) with a cluster of multiple bound sperm (scored as >10), and a *bncr*^−/−^ egg (bottom) with no bound sperm. E – Quantification of sperm-egg binding. Eggs were classified as either >10 sperm bound or <10 sperm bound. Mean and standard deviation are indicated (p < 0.002, Mann-Whitney test; n = total number of eggs; N = number of biological replicates).

Successful fertilization is a multistep process. A role for Bouncer at the egg membrane implies that it could promote the approach of sperm to the egg, or sperm-egg binding/fusion. To assess whether Bouncer influences sperm recruitment, we used live imaging to determine if MitoTracker-labeled wild-type sperm can approach the micropyle of Bouncer-deficient eggs. We found that multiple sperm were recruited to the micropyle area of Bouncer-deficient eggs within seconds after sperm addition, similar to wild-type eggs (Fig. 3C; Supplementary Movie 2). Thus, Bouncer is not providing an essential attractive cue that guides sperm towards the egg/micropyle.

During live imaging, multiple sperm were recruited into the narrow opening of the micropyle simultaneously, rendering a more detailed analysis of the sperm-egg binding capability unfeasible. To evaluate a potential role for Bouncer in sperm-egg binding, we therefore chose a different experimental setup. Wild-type and Bouncer-deficient eggs were activated by adding spawning medium in the absence of sperm. After chorion elevation, the eggs were manually dechorionated to expose the entire egg surface, incubated with MitoTracker-labeled sperm, gently washed and examined by confocal microscopy. In the case of wild-type eggs, we found that large clusters of sperm (>10 sperm) remained bound to the egg membrane (Fig. 3D, E). In contrast, only a few individual sperm (<10) remained attached to most Bouncer-deficient eggs (p < 0.002, Mann-Whitney test) (Fig. 3D, E). These results suggest that Bouncer promotes stable sperm-egg binding.

The presence of Bouncer homologs among vertebrates, yet high degree of amino acid sequence divergence particularly among fish homologs (Fig. 1C; Supplementary Fig. 2A, B), raised the interesting possibility that Bouncer might contribute to the species-specificity of fertilization. To test this hypothesis, we first evaluated whether Bouncer from medaka (*Oryzias latipes*) could rescue the fertility defect of zebrafish *bncr*^−/−^ females. Medaka was chosen because of its large evolutionary distance from zebrafish (>150 million years), its inability to cross-hybridize with zebrafish, and the low (40%) level of amino acid identity between mature zebrafish and medaka Bouncer proteins (Fig. 4A). We generated *bncr*^−/−^ zebrafish that ubiquitously express medaka Bouncer (*bncr*^−/−^; tg[*ubi:medaka-bncr*], called in brief “transgenic medaka Bouncer” fish). We crossed wild-type male zebrafish with *bncr*^−/−^ females, transgenic medaka Bouncer females, and zebrafish Bouncer females. Transgenic medaka Bouncer females displayed a substantially decreased fertilization rate (average rate of 0.62%) compared to zebrafish Bouncer females (average rate of 78.6%), similar to the fertilization rate of *bncr*^−/−^ females (average rate of 0.11%) (Fig. 4B). Thus, medaka Bouncer does not efficiently rescue the fertilization defect of *bncr*^−/−^ zebrafish.

**Figure 4:**
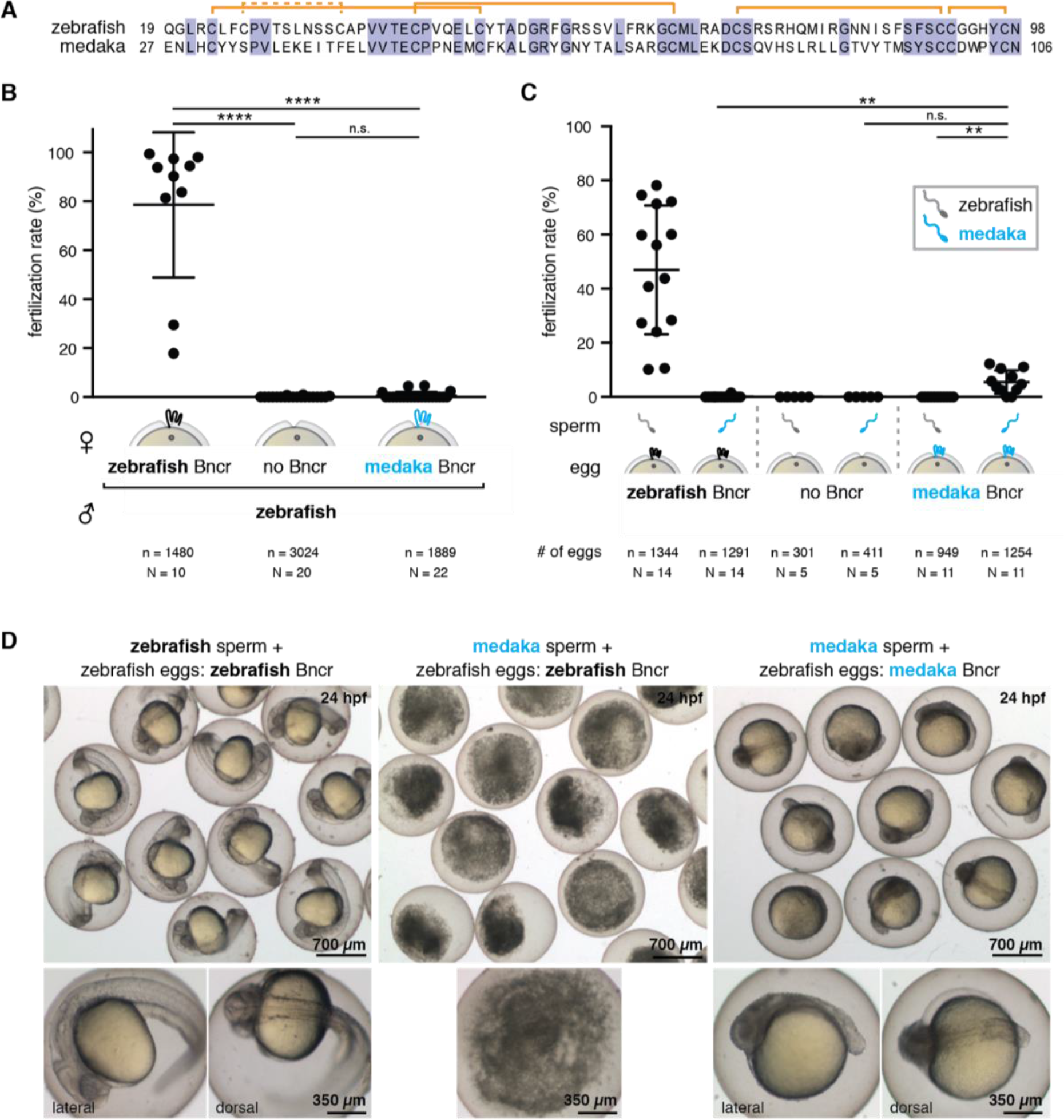
Bouncer mediates species-specific fertilization. A – Mature zebrafish and medaka Bouncer have only 40% amino acid sequence identity. Shown is a protein sequence alignment of the two mature proteins. Orange brackets indicate predicted disulfide bonds. B – Medaka Bouncer does not efficiently rescue the fertilization defect of zebrafish *bncr*^−/−^ females. Expression of medaka Bouncer in a zebrafish *bncr*^−/−^ background results in similar fertilization rates by zebrafish sperm than in the case of *bncr*^−/−^ mutant eggs. Fertilization rates are much lower in comparison to wild-type eggs expressing zebrafish Bouncer. Data is shown for all (17) medaka Bouncer-expressing females tested. Mean and standard deviation areindicated (Kruskal-Wallis test with Dunn’s multiple comparisons test: wild-type × wild-type vs. *bncr*^−/−^ × wild-type, adj. p < 0.0001; vs. medaka Bouncer × wild-type, adj. p < 0.0001; *bncr*^−/−^ × wild-type vs. medaka Bouncer × wild-type, adj. p > 0.9999, n.s., n = total number of eggs; N = number of biological replicates). C – Medaka Bouncer is sufficient to allow entry of medaka sperm into zebrafish eggs. While medaka sperm was unable to enter wild-type zebrafish eggs, *in vitro* fertilization of zebrafish *bncr*^−/−^ eggs expressing medaka Bouncer with medaka sperm resulted in an average fertilization rate of 5.5%. Data is shown only for the subset of medaka Bouncer-expressing females (4 out of 17) that showed any fertilization ability (data for all 17 females is shown in Supplementary Fig. 5A). Mean and standard deviation are indicated (Kruskal-Wallis test with Dunn’s multiple comparisons test: medaka sperm on zebrafish vs. medaka Bouncer eggs, adj. p = 0.0096; zebrafish sperm on medaka Bouncer eggs vs. medaka sperm on medaka Bouncer eggs, adj. p = 0.0085; medaka sperm on *bncr*^−/−^ zebrafish eggs vs. medaka Bouncer eggs, adj. p = 0.0549, n.s.; n = total number of eggs; N = number of biological replicates). D – Fertilization of zebrafish eggs expressing only medaka Bouncer yields medaka-zebrafish hybrid embryos that are viable for up to two days. Left: Wild-type zebrafish embryos fertilized by zebrafish sperm. Middle: Incubation of wild-type zebrafish embryos with medaka sperm does not result in fertilization and yields decomposed eggs after 24 hours. Right: Zebrafish eggs expressing only medaka Bouncer were fertilized by medaka sperm and develop into hybrid embryos (Supplementary Fig. 5B) with anterior-posterior axis formation after 24 hours. All images were taken at 24 hours post-fertilization (hpf).

Our finding that transgenic medaka Bouncer eggs were not efficiently fertilized by zebrafish sperm supported our hypothesis that Bouncer might influence species-specific gamete interactions. To directly test this possibility, we performed a series of IVF experiments. As expected, wild-type zebrafish eggs exhibited high fertilization rates with zebrafish sperm (average rate of 49.0%), but were not fertilized by medaka sperm (Fig 4C). Moreover, zebrafish *bncr*^−/−^ eggs were fertilized by neither sperm (Fig 4C). Remarkably, eggs from transgenic medaka Bouncer females were fertilized by medaka but not zebrafish sperm. The average fertilization rate of all transgenic medaka Bouncer females (*17*) tested was 2.4% (Supplementary Fig. 5A). While 13 out of 17 females were completely infertile, eggs from the remaining four females were fertilized by medaka sperm at an average rate of 5.5% (Fig. 4C). The observed difference in fertility rates is likely due to independent insertions of the medaka Bouncer transgene. The resulting embryos were medaka-zebrafish hybrids and not haploid zebrafish embryos (Supplementary Fig. 5B). Hybrid embryos underwent cleavage cycles and gastrulation movements (Supplementary Fig. 5C), and displayed anterior-posterior axis formation after 24 hours (Fig 4D), but did not survive past 48 hours. These results demonstrate that Bouncer is necessary and sufficient for mediating species-specific fertilization in fish.

## Discussion

In this study, we discovered that the Ly6/uPAR protein Bouncer is a key factor in vertebrate fertilization. Zebrafish Bouncer localizes to the oocyte membrane and is required for female fertility by enabling sperm entry. Intriguingly, we find that Bouncer is not only required for efficient sperm-egg binding, but also sufficient to enable cross-species fertilization. Thus, this is the first report of a protein in any organism allowing entry of another species’ sperm. In light of this crucial function, we chose the name “Bouncer” in reference to the colloquial name for a doorman at a bar. Like a security guard, Bouncer is decisive for allowing conspecific sperm to enter, while keeping heterospecific sperm out.

Thus far, the only known interacting membrane-bound proteins on vertebrate sperm and egg are IZUMO1 and JUNO in mammals (*1–3*, *7*), yet these two proteins do not confer species-specificity to fertilization (Bianchi and Wright 2015). Bouncer is the first protein known to function in sperm-egg binding in a non-mammalian vertebrate, and it is the first membrane-anchored protein known to mediate species-specificity of fertilization in any vertebrate. Our finding that ectopic expression of another species’ Bouncer is sufficient to allow cross-species fertilization strongly suggests that Bouncer has a direct, species-specific interaction partner on sperm. The future identification of this interaction partner will be a crucial next step to unravel the mechanism behind Bouncer’s function.

In many organisms, species-specificity of fertilization is mediated between sperm-membrane bound proteins and proteins localized to the egg coat (*15*, *16*). For example, in sea urchin, the vitelline envelope protein EBR1 (egg bindin receptor protein-1) binds specifically to the sperm membrane protein Bindin (*17–19*). Similarly, the egg coat protein Verl of abalone is species-specifically bound by Lysin, a small protein that is secreted by sperm (*20–23*). While Lysin has no known homolog in vertebrates, Verl shows structural homology to the mammalian zona pellucida protein ZP2 (*22*), which was shown to be involved in species-specific binding of the sperm to the zona pellucida in mouse and humans (*24*). Thus, Bouncer is the first known protein to mediate species-specific binding of sperm not to an egg coat, but directly to the egg membrane.

Bouncer and SPACA4, both members of the large Ly6/uPAR superfamily, have opposing germline-specific expression patterns in externally versus internally fertilizing organisms. Although it is not clear why different vertebrates show opposing expression patterns, one can speculate that in externally fertilizing species, the oocyte-specific expression of Bouncer could contribute to post-copulation female mate choice (also called cryptic female mate choice) (*25*). Vertebrates performing external fertilization cannot guarantee that only conspecific sperm reaches the egg by pre-mating choice (*26*, *27*). Oocyte-expressed proteins like Bouncer could therefore support the selection of conspecific sperm. Our work on Bouncer also raises the intriguing possibility that SPACA4 might play a similarly fundamental role in mammalian fertilization, albeit from the side of the male. While a knock-out study for *Spaca4* in mice has not yet been reported, this idea is consistent with the localization of SPACA4 to the inner acrosomal membrane of sperm and the observed antibody-blocking effect *in vitro* (*14*). Future experiments that address the *in vivo* function of mammalian SPACA4 during fertilization will be of interest. Given that both genes are restricted to the germline, our findings in fish might have direct relevance for fertilization in mammals.

## Acknowledgements

We thank Kristin Tessmar-Raible and Bruno M. Fontinha for generously providing medaka fish and expertise, Alex Schier for his generous support during the start of this project and for valuable feedback on the manuscript, and Jamie Gagnon for his generous help in generating the *bouncer* mutant and obtaining germline RNA-Seq data. We also thank Maria Novatchoka and Luis Enrique Cabrera Quio for help with RNA-Seq data mapping and gene expression analyses, Karin Panser for help with genotyping, the IMP animal facility personnel, especially Julia König and Friedrich Ecker, for their excellent care of our fish, and Thomas Heuser and Nicole Fellner from the VBCF EM facility for help with EM. We furthermore acknowledge the following people for providing reagents: Mathias Madalinski for synthesizing Bouncer peptides for antibody production, Jeff Farrell for providing the sfGFP plasmid, and Carl-Philipp Heisenberg for providing the tg[*lyn-tdTomato*] fish line. We would also like to thank Angela Anderson (Life Science Editors), Alex Stark, Carl-Philipp Heisenberg, Luisa Cochella, Masahito Ikawa and Yoshitaka Fujihara for helpful comments on the manuscript.

## Author contributions

S.H. and A.P. conceived the study. S.H. performed most experiments except experiments regarding species-specificity, which were performed by K.R.G., and generation of RNA-sequencing data and bncr^−/−^ fish, which were performed by A.P. S.H., K.R.G. and A.P. analyzed the data. A.S. performed the phylogenetic analysis. S.H. and A.P. wrote the manuscript with input from K.R.G. and A.S.

## Funding

This work was supported by the Research Institute of Molecular Pathology (IMP), Boehringer Ingelheim and the Austrian Academy of Sciences. S.H. is supported by a DOC Fellowship from the Austrian Academy of Sciences. A.P. is supported by the HFSP Career Development Award (CDA00066/2015) and the FWF START program.

## Supplementary Materials

### Materials and methods

#### Fish lines and husbandry

Zebrafish (*Danio rerio*) were raised according to standard protocols (28°C water temperature; 14/10 hour light/dark cycle). TLAB fish, generated by crossing zebrafish AB and the natural variant TL (Tübingen/Tüpfel Longfin) stocks, served as wild-type zebrafish for all experiments. *Bouncer* mutant zebrafish and transgenic lines were generated as part of this study and are described in detail below. The zebrafish line tg[*actb2:lyn-tdTomato*] was kindly provided by the Heisenberg lab and used for membrane labeling. Wild-type medaka fish (*Oryzias latipes*, strain CAB) were raised according to standard protocols (28°C water temperature; 16/8 hour light/dark cycle) in the fish facility of the Max F. Perutz Laboratories Vienna, and were kindly provided by the Tessmar-Raible lab.

All fish experiments were conducted according to Austrian and European guidelines for animal research and approved by local Austrian authorities (animal protocol BMGF-76110/0017-II/B/16c/2017 for work with zebrafish; animal protocols BMWFW-66.006/0012-WF/II/3b/2014 and BMWFW-66.006/0003-WF/V/3b/2016 for work with medaka).

#### RNA-Seq from the zebrafish germline

Testes, ovaries, and mature oocytes (obtained by squeezing) of wild-type (TLAB) zebrafish were collected (two biological replicates for each sample). Total RNA was isolated using the standard TRIzol (Invitrogen, Waltham, Massachusetts) protocol, and genomic DNA was removed by TURBO DNase treatment followed by phenol/chloroform extraction. Strand-specific libraries for 76-bp paired-end sequencing were prepared from cDNA by the Broad Institute Sequencing Platform. Libraries were sequenced on the Illumina HiSeq 2000. Reads were aligned to GRCz10, using the Ensembl transcriptome release 88. A custom file was generated by adding *bouncer* based on its position coordinates (exon = chr18:50858259-50858859 (- strand); CDS = chr18:50858285-50858663 (- strand)). The following command was used to map each sample: ‘tophat - o <output directory> −p 16 --library-type fr-firststrand-no-novel-juncs-g 1 -G <Custom_gene_table> <Bowtie2_genome_index> <fastq_reads>”. Quantification of transcript levels was determined using cuffnorm with the following command “cuffnorm −p 22 --library-type=fr-firststrand -L < labels > −o <output_directory> <Custom_gene_table> <aligned_reads.bam file>. Coverage tracks were displayed in the Integrative Genomics Viewer (IGV) (http://software.broadinstitute.org/software/igv/). The RNA-Seq data set was deposited to Gene Expression Omnibus (GEO) and is available under GEO acquisition number GSE111882.

#### Protein domain analysis of Bouncer

Protein domains were assigned using hmmscan (v. 3.1b2) (*33*) and profile hidden Markov models derived from PFAM (v. 31.0, March 2017) (*34*), with a significant E-value threshold of 0.01. The domain search classified zebrafish Bouncer as a member of the Ly6/uPAR protein family (Pfam domain UPAR_LY6_2, E-value 1.4e-05). The conserved domain covers nearly the entire protein (residues 17-100), apart from the amino-terminal signal peptide (predicted signal peptide cleavage site after amino acid 18 (VLP-QG) based on SignalP 4.1 (http://www.cbs.dtu.dk/services/SignalP/) (*35*)) and a putative transmembrane region at the carboxy-terminus (amino acids 109-125, based on TMpred (https://embnet.vital-it.ch/software/TMPRED_form.html) (*36*)). Based on GPI-prediction tools (http://mendel.imp.ac.at/gpi/gpi_server.html) (*37*), zebrafish Bouncer is a GPI-anchored protein (GPI-anchor site: N98; p-value 2.1e-03). Glycosylation site prediction tools (http://www.cbs.dtu.dk/services/NetNGlyc/ and http://www.cbs.dtu.dk/services/NetOGlyc/) identified two possible N-glycosylation sites in zebrafish Bouncer: amino acids N32 and N84.

#### Homolog identification and phylogenetic analysis

An NCBI-BLASTP search (version 2.6.0+) (*38*) with *Danio rerio* Bouncer (NCBI gene *LOC101885477*; XP 005173770.1; gene ID:101885477) against the human proteome (NCBI version 12/2017, 357531 non-redundant entries) hit only to the SPACA4 protein (E-value 9.72e-06). A reciprocal search with human SPACA4 protein against the zebrafish proteome (60215 non-redundant entries) returned Bouncer as the best hit (E-value 1.43e-04).

An NCBI-BLASTP search with *Danio rerio* Bouncer against the NCBI non-redundant (nr) protein database (version 11/2017, 138830414 entries) resulted in highly significant hits (below E-value 1e-10) to proteins from bony fishes such as *Oncorhynchus mykiss* (rainbow trout), *Hippocampus comes* (tiger tail seahorse) and *Oreochromis niloticus* (Nile tilapia). The closest hits outside of bony fish were to SPACA4 (sperm acrosome membrane-associated protein 4) family proteins in *Myotis brandtii* (Brandt’s bat, E-value 1.20e-09) and *Xenopus tropicalis* (tropical clawed frog, E-value 5.09e-09). Since no medaka (*Oryzias latipes*) Bouncer could be detected in the NCBI nr database, we performed a TBLASTN search within the NCBI EST division and identified the cDNA clone McF0051J12-MGRbd1 (NCBI EST accession AM154129) as best hit (E-value 3e-15), coding for a 131 amino acid gene product. The homolog relationship was confirmed in a reciprocal BLAST search against the zebrafish genome and in a phylogenetic tree with selected members of the Ly6/uPAR family (see below).

#### Phylogenetic tree

Starting with *Danio rerio* Bouncer, we collected all significant (E-value 0.001) hits in an NCBI-BLASTP search within the non-redundant protein database. To extend the Ly6/uPAR family, these hits were used as a query in an additional search step applying significant criteria (E-value 0.001) for hit collection. Forty-two species were selected to represent a wide taxonomic range. Within each species, the sequences were reduced to 80% identity with cd-hit (*39*). Proteins were aligned with mafft (−linsi v7.313) (*40*), and sequences with long gaps were removed. The medaka (*Oryzias latipes*) Bouncer sequences, and one Ly6/uPAR protein of the shark *Squalus acanthias* were added (see Supplemental Table 1), realigned, resulting in a final set of 192 full-length sequences. The respective coding sequences were extracted from the NCBI website (where available, see Supplementary Table), and aligned with RevTrans using the protein sequence alignment (v. 1.4) (*41*). Aligned columns with less than 50% sequence information were removed. The alignment was furthermore restricted to the region corresponding to the putative mature peptide of *Danio rerio* Bouncer using Jalview (*42*). A maximum likelihood phylogenetic tree was calculated with IQ-TREE (v. 1.5.5) (*43*) with codon model selection using ModelFinder (*44*) and 1000 ultrafast bootstrap replicates (*45*). The tree was visualized in iTOL (*46*).

#### Generation of *bouncer* knockout fish

*Bouncer* mutant fish were generated by Cas9-mediated mutagenesis, targeting the predicted (*9*), un-annotated single-exon gene downstream of the *ppfia1* gene (location of the *bouncer* ORF in GRCz10: chr18:50,858,463-50,858,837). A guide RNA (sgRNA) targeting the region encoding the N-terminal signal peptide of Bouncer was generated according to published protocols (*47*) by oligo annealing followed by T7 polymerase-driven *in vitro* transcription (gene-specific targeting oligo: bouncer_gRNA; common tracer oligo). Cas9 protein and *bouncer* sgRNA were co-injected into the cell of one-cell stage TLAB embryos. Putative founder fish were outcrossed to TLAB wild-type fish. A founder fish carrying a germline mutation in the *bouncer* gene was identified by a size difference in the *bouncer* PCR amplicon in a pool of embryo progeny (primer: bouncer_gt_F and bouncer_gt_R). Embryos from this founder fish were raised to adulthood. Amplicon sequencing of adult fin-clips (primers: bouncer_gt_F and bouncer_gt_R) identified the mutation, namely a 13-nt deletion, which introduces a premature termination codon at the end of the Bouncer signal peptide sequence.

Genotyping of *bouncer* mutant fish was performed by PCR (primers: bouncer_gt_F and bouncer_gt_R). Detection of the 13-nt deletion was performed by standard gel electrophoresis using a high-percentage agarose gel either a) without further processing of the PCR product, or b) after EcoNI digest, which cuts only the wild-type but not the mutant amplicon (wild-type amplicon: 234-nt, 13-nt deletion amplicon: 221-nt, EcoNI-digested wild-type amplicon: 126-nt + 108-nt).

Homozygous *bouncer*^−/−^ knockout fish (*bncr*^−/−^) were generated by incrossing *bncr*^+/−^ fish or by crossing a *bncr*^−/−^ male to a *bncr*^+/−^ female. Alternatively, *bncr*^−/−^ fish were generated by crossing a *bncr*^−/−^ female, carrying a sfGFP-tagged or untagged *bouncer* rescue construct (*tg[ubi:sfGFP-bncr, cmlc2:eGFP]* or *tg[ubi:bncr, cmlc2:eGFP]*), to a *bncr*^−/−^ male and screening for the absence of green fluorescent hearts.

#### Generation of zebrafish expressing transgenic Bouncer

The predicted *bouncer* transcript unit, including its 24-nt 5’UTR, 378-nt ORF encoding the 125-amino acid zebrafish Bouncer protein, and 236 nts of its 3’UTR, was amplified by PCR from cDNA derived from 2-4 cell stage zebrafish embryos (primers: bouncer_F; bouncer_R) and cloned into the pSC vector (Strataclone) to generate *pSC-bncr*. sfGFP (superfold GFP)-tagged Bouncer (*pSC-signalpeptide-sfGFP-bncr*) was generated by digesting *pSC-bncr* with EcoNI, which cuts *bncr* at the end of the predicted signal peptide sequence (…VVLP-Q…), and by inserting a sfGFP-encoding PCR fragment (primers: sfGFP-bouncer_F; sfGFP-bouncer_R) in frame immediately after the predicted signal peptide sequence (…WLP-sfGFP…) at the N-terminus of the mature Bouncer protein by Gibson cloning (*48*). Untagged *bouncer* and *sfGFP-tagged bouncer* were subsequently amplified by PCR and subcloned into vectors for Tol2-based transgenesis (BamHI/NheI-cut *pTol2-ubi:MCS, cmlc2:GFP* or BamHI/AgeI-cut *pMTB-actb2:H2B-Cerulean* (kind gift from Sean Megason)) to generate *pTol2-ubi:bncr, cmlc2:eGFP; pTol2-ubi:sfGFP-bncr, cmlc2:eGFP;* and *pMTB-actb2:sfGFP-bncr*.

To generate sfGFP-tagged Bouncer with mutations of the two predicted N-glycosylation sites (N32, N84), a gBlock (SH043; IDT) encoding part of the sfGFP ORF and the N32A/N84A (AAC -> GCC) mutated *bouncer* ORF was cloned into ClaI/XmaI-digested *pTol2-ubi:sfGFP-bncr*, *cmlc2:eGFP* to obtain *pTol2-ubi:sfGFP-bncr*^*N32A, N84A*^, *cmlc2:eGFP*.

To generate secreted sfGFP-tagged Bouncer lacking the GPI-anchor site and C-terminal transmembrane domain (sfGFP-Bouncer^noTM^), a gBlock (SH044, IDT) encoding the truncated *bouncer* ORF was cloned into ClaI/XmaI-digested *pTol2-ubi:sfGFP-bncr*, *cmlc2:eGFP* to obtain *pTol2-ubi:sfGFP-bncr*^*noTM*^, *cmlc2:eGFP*.

To generate transgenic zebrafish lines expressing medaka Bouncer, medaka Bouncer (NCBI nucleotide accession number AM154129.1) was PCR-amplified from cDNA derived from oocyte mRNA (primers: medaka_bouncer_F and medaka_bouncer_R) and cloned into a vector for Tol2-based transgenesis (BamHI/NheI-cut *pTol2-ubi:MCS*, *cmlc2:eGFP*) *to obtain p Tol2-ubi:medaka-bncr*, *cmlc2:e GFP*.

Zebrafish lines expressing transgenic Bouncer were generated by injecting *bouncer* expression constructs with *Tol2* mRNA into TLAB or *bncr*^+/−^ zebrafish embryos (4 ng/μl of medaka Bouncer plasmid or 15 ng/μl of all other purified plasmids in RNase-free water, 9.3 ng/μl for medaka Bouncer transgenesis or 35 ng/μl *Tol2* mRNA for all others, 0.083% phenol red solution (Sigma-Aldrich)), following standard procedures. Injected embryos with high expression of the fluorescent marker (either *cmlc2:eGFP*, or *actb2:sfGFP-bncr*) at one day post fertilization were raised to adulthood. Putative founder fish were crossed to *bncr*^−/−^ or *bncr*^+/−^ fish, and the progeny was screened after 48 hours for the expression of the fluorescent marker. GFP-positive embryos were then raised to adulthood and genotyped for *bouncer*. Functionality of the expression constructs was determined by assessing the ability to rescue the near-sterility of *bncr*^−/−^ females.

The following transgenic zebrafish lines were generated (in wild-type (TLAB), *bncr*^+/−^ or *bncr*^−/−^ backgrounds):

- *tg[ubi:bncr, cmlc2:eGFP]*
- *tg[ubi:sfGFP-bncr, cmlc2:eGFP]*
- *tg[actb2:sfGFP-bncr]*
- *tg[ubi:sfGFP-bncr*^*noTM*^, *cmlc2:eGFP]*
- *tg[ubi:sfGFP-bncr*^*N32A, N84A*^, *cmlc2:eGFP]*
- *tg[ubi:medaka-bncr, cmlc2:eGFP]*

The following expression constructs were determined to be fully functional (rescuing the near-sterility *of bncr*^−/−^ females: *ubi:bncr*, *ubi:sfGFP-bncr*, *actb2:sfGFP-bncr*, and *ubi:sfGFP-bncr*^*N32A, N84A*^.

#### Imaging of live embryos

Live embryos and larvae were imaged in their chorions in E3 medium (5 mM NaCl, 0.17 mM KCl, 0.33 mM CaCl_2_, 0.33 mM MgSO_4_, 10^−5^ % Methylene Blue) using a Stereo Lumar.V12 fluorescence microscope (Zeiss).

#### Antibodies

To generate an antibody specific for zebrafish Bouncer, two *in vitro*-synthesized peptides (CYADGRFGRSSVLFRKG and CSRSRHQMIRGNNIS) were used for immunization of rabbits (Eurofin). The final bleed was purified against each of the two peptides separately. For this purpose, each peptide was bound to a column by first dissolving 5 mg of the peptide in 3 ml of 20 mM Hepes and 1 mM EDTA. The peptide solution was then mixed with 300-310 mg maleimide-activated POROS and flushed with argon. Afterwards, the suspension was incubated at RT for 1 hour with gentle mixing. The column (4.6 × 50 mm) was packed using column packer (Applied Biosystems, buffer: HBS; flow: 10 ml/min for 20-25 min) on HPLC. For the affinity purification (monitoring at 280 nm, flow: 7.5 ml/min), 5 ml of the serum were injected onto the column, followed by a wash with HBS. For the first elution MgCl_2_-containing buffer (1.5 M MgCl_2_, 50 mM NaAc, pH 5.2), and for the second elution glycine-buffer (0.1 M glycine, 0.1 M NaCl pH 2.45) was used. The purified antibody was dialyzed using HBS buffer and stored in 10% glycerol at 4°C. The different antibody fractions were tested for their reactivity against Bouncer by western blotting. The glycine eluate purified against peptide CYADGRFGRSSVLFRKG gave the strongest and cleanest signal for Bouncer and was therefore used for all experiments.

#### Analysis of zebrafish egg and embryo lysates by western blotting

For western blotting of zebrafish egg and embryo lysates, freshly laid eggs or dome stage embryos (as indicated) were dechorionated manually in E3 medium (5 mM NaCl, 0.17 mM KCl, 0.33 mM CaCl_2_, 0.33 mM MgSO_4_, 10^−5^% Methylene Blue). Afterwards, the yolk was manually removed using forceps, and the egg/embryo caps were frozen in liquid nitrogen and stored at −80°C until usage. Western blotting was performed following standard protocols. In brief, the equivalent of 20 eggs (or embryos) were loaded per lane. SDS-PAGE was performed using Mini-PROTEAN^®^ TGX™ Precast Protein Gels or 10-20% Mini-PROTEAN^®^ Tris-Tricine Gels. Blotting was performed using a wet-blot system (BioRad). The following antibodies were used in western blotting: anti-Bouncer, rabbit, 1:500 (generated as described above), antitubulin (mouse, Sigma-Aldrich T6074, 1:20000).

#### Analysis of glycosylation of Bouncer

Manually deyolked egg and embryo samples were deglycosylated overnight under nondenaturing conditions using Protein Deglycosylation Mix (New England Biolabs) according to the manufacturer’s protocol. The glycosylation state of endogenous Bouncer was assessed by western blotting using Bouncer-specific antibodies.

#### *In situ* hybridization

*Bouncer* DIG-labeled RNAs (sense and antisense) were generated by digesting *pSC-bncr* with EcoRV (antisense) or BamHI (sense), followed by *in vitro*-transcribing the linearized plasmids using T7 (antisense) or T3 (sense) polymerases (Roche) and DIG RNA labeling mix (Roche). *In situ* hybridization in zebrafish embryos was performed according to standard protocols, using *anti-bouncer* or *sense-bouncer* (negative control for nonspecific staining) DIG-labeled RNA probe. BCIP/NBT/alkaline-phosphatase-stained embryos were dehydrated in methanol and imaged in BB/BA using a Stereo Lumar.V12 fluorescence microscope (Zeiss) and an Axioplan 2 microscope (Zeiss) equipped with a DFC320 camera (Leica).

#### *In vivo* fertilization in fish

The evening prior to mating, male and female fish were separated in breeding cages (one male and one female per cage). In the morning of the next day, male and female fish were allowed to mate by removing separators. Eggs were collected from the bottom of the breeding cages and kept at 28°C in E3 medium. The rate of fertilization was assessed about 3 hours post laying. By this time, fertilized embryos have developed to ~1000 cell stage embryos, while unfertilized embryos resemble 1-cell stage embryos.

#### *In vitro* fertilization in fish

*In vitro* fertilization experiments were performed following standard procedures (*49, 50*). The evening prior to the zebrafish egg and sperm collections, male and female zebrafish were separated in breeding cages (one male and one female per cage).

To collect mature, un-activated eggs, female zebrafish were anesthetized using 0.1% Tricaine (25× stock solution in dH_2_O, buffered to pH 7-7.5 with 1 M Tris pH 9.0). After being gently dried on a paper towel, the female was transferred to a dry petri dish, and eggs were carefully expelled from the female by applying mild pressure on the fish belly with a finger and stroking from anterior to posterior. The eggs were separated from the female using flat forceps or a small paintbrush, and the female was transferred back to the breeding cage filled with fish water for recovery.

To collect un-activated zebrafish sperm, male zebrafish were anesthetized using 0.1% Tricaine. After being gently dried on a paper towel, the male fish was placed belly-up in a slit in a damp sponge under a stereomicroscope with a light source from above. Sperm was collected into a glass capillary by mild suction while gentle pressure was applied to the fish belly using flat forceps. The sperm of several males (2–5) was stored in ice-cold Hank’s saline (0.137 M NaCl, 5.4 mM KCl, 0.25 mM Na_2_HPO_4_, 1.3 mM CaCl_2_, 1.0 mM MgSO_4_, 4.2 mM NaHCO_3_) on ice. The male was transferred back to the breeding cage filled with fish water for recovery. To label sperm with MitoTracker, sperm was incubated in 0.5 μM MitoTracker™ Deep Red FM (Thermo Fisher Scientific) on ice for >10 minutes prior to *in vitro* fertilization. Un-activated medaka sperm was collected according to published procedures (*50*). The sperm was stored in ice-cold Hank’s saline (0.137 M NaCl, 5.4 mM KCl, 0.25 mM Na_2_HPO_4_, 1.3 mM CaCl_2_, 1.0 mM MgSO_4_, 4.2 mM NaHCO_3_) on ice until usage.

To fertilize eggs *in vitro*, an aliquot of 25-50 μl unlabeled or MitoTracker-labeled sperm solution was added to eggs of potentially different genotypes (e.g. collected from wild-type, bncr^−/−^ or transgenic females) in individual petri dishes. Sperm addition was immediately followed by adding 500 μl-1 ml of E3 medium to simultaneously activate sperm and eggs. After 2 minutes, petri dishes were filled with E3 medium and kept at 28°C. The rate of fertilization was assessed about 3 hours post sperm addition. By this time, fertilized embryos have developed to ~1000-cell stage embryos, while unfertilized embryos resemble 1-cell stage embryos.

#### Analysis of oocyte activation and polar body extrusion

Oocyte activation was assessed by monitoring chorion elevation after addition of E3 medium to un-activated eggs that had been obtained by squeezing.

To assess polar body extrusion, eggs were *in vitro* fertilized and fixed after 3, 10 and 20 minutes in 3.7% formaldehyde in PBS at 4°C overnight. Embryos were washed in PBST, permeabilized in Methanol and kept at −20°C. For immunostaining, embryos were transferred back to PBST, dechorionated manually using forceps, and stained with a rabbit anti-*γ*-tubulin antibody (T3559 Sigma, used at 1:1000; secondary antibody: goat anti-rabbit Alexa488 (A-11034 Thermo Scientific, used at 1:250)) and 1× DAPI in PBS. Embryos were mounted in 1% low-melt agarose on glass-bottom dishes (Ibidis) and imaged at an inverted LSM880 confocal microscope (Zeiss) with a 20× objective lens at 2× magnification.

#### Analysis of cytoplasmic streaming

Embryos derived from *in vivo* mating crosses between wild type males and *wild-type* or *bncr*^−/−^ females were collected immediately after being laid and dechorionated with pronase (1 mg/ml). Dechorionated embryos were injected with 1 nl of red fluorescent FluoSpheres^®^ NeutrAvidin^®^ labeled microspheres (ThermoFisher, 1:5 dilution in 0.083% phenol red) in the lower third of the yolk. Directly after injection, embryos were mounted laterally in glass-bottom dishes (Ibidi) in 0.8% LM (low-melt) agarose in PBS. Embryos were imaged using an inverted LSM 780 confocal microscope (Zeiss) with a 10× objective lens at 1× magnification for 200 cycles over 3 hours. Embryos were kept at 28°C during imaging.

Statistical analysis was performed using Prism software (Graphpad). Small accumulations of beads were tracked (on average, 8 bead accumulations per embryo) using the manual tracking function in Fiji (*51*), which recorded the velocity for each time point. The mean velocity that each bead traveled in each embryo was calculated; these values were then averaged to produce the total mean velocity of all beads in a single embryo. An unpaired t-test was used to compare the total mean velocities between wild type and *bouncer* knockout embryos.

#### Scanning Electron microscopy

Eggs from wild-type and bncr^−/−^ females were squeezed into dry petri dishes. E3 medium was added for activation, and after 10 seconds to 1 minute, glutaraldehyde (2.5%) was added for initial fixation. The samples were then stored at 4°C overnight. The eggs were transferred to 2.5% glutaraldehyde in 0.1 mol/l sodium phosphate buffer, pH 7.4 and incubated for 2 hours. Eggs were rinsed with the same buffer and post-fixed in 1% osmium tetroxide in ddH_2_O. After rinsing with ddH_2_O samples were dehydrated by a 10-minute treatment in dimethoxipropane followed by 3 incubation steps in anhydrous acetone for 30 minutes each. The eggs were then treated with a 1:1 mixture of anhydrous acetone and hexamethyldisilazane (HMDS) followed by 3 HMDS treatments for one hour each. After air-drying for several hours in a fume hood, eggs were gold-sputter coated and examined in a Hitachi TM-1000 tabletop scanning electron microscope operated at 15 kV and equipped with a highly sensitive semiconductor backscattered electron detector.

#### Coomassie staining of the micropyle

To stain the micropylar region of the chorion, eggs from normal *in vivo* crosses were collected ~10 minutes after egg laying and fixed in 3.7% formaldehyde in PBS at 4°C overnight. Embryos were washed in PBST and stained with Coomassie Brilliant Blue G (final concentration: 0.2% in 10% DMSO in PBS) for 10 minutes at room temperature. Embryos were rinsed in PBS. The chorion was manually separated from the embryo using forceps, and flat-mounted on a glass slide. Images of the micropylar region were taken with bright field optics using an Axio Imager.Z1 microscope (Zeiss) with 20× or 40× objectives. Quantification of the diameter of the micropylar region was performed in FIJI, using the measurement tool. The average diameter of each micropyle was calculated from two perpendicular measurements of each micropylar opening. Statistical analyses were performed using Prism software (Graphpad). Unpaired t-tests were used to compare the average diameters of the micropyles between wild-type and *bouncer* knockout embryos.

#### Intra-cytoplasmic sperm injection (ICSI) in zebrafish

Sperm of two males was pooled in 50 μl of Hank’s saline (0.137 M NaCl, 5.4 mM KCl, 0.25 mM Na_2_HPO_4_, 1.3 mM CaCl_2_, 1.0 mM MgSO_4_, 4.2 mM NaHCO_3_). To assess the sperm concentration, a small aliquot of sperm was diluted 1:100 in Hank’s saline, and the sperm number was counted using a cell counting chamber (Neubauer pattern, 0.1 mm depth, 0.0025 m^2^, BRAND^®^) on a cell culture microscope (Axio, Vert.A1, Zeiss). The sperm solution was diluted to a working concentration of 2000-4000 sperm/μl in 5% polyvinylpyrrolidone (PVP) in E3 medium and filled into the injection needle using Microloader™ pipette tips (Eppendorf). The sperm number was checked by injecting 0.5 nl of the sperm mix into seven 2 μl droplets of a 1× DAPI solution containing 10 mg/ml digitonin in 5% PVP, using a standard microinjection apparatus (PV820 Pneumatic PicoPump, World Precision Instruments). The sperm in the droplets was imaged using a cell culture microscope (Axio, Vert.A1, Zeiss). If the sperm number was correct (1-2 sperm/0.5 nl), un-activated, mature eggs were collected from a female and kept in Hank’s saline (+ 0.5% BSA) during injection to slow down egg activation. During injection, eggs were held in place in wedge-shaped wells of a 1.5% agarose petri dish with the help of flat forceps. Sperm was injected into the cell as close as possible to the micropyle. After injection, the eggs were immediately transferred to E3 medium and kept at 28°C. After 3-5 hours, the injected eggs were analyzed for cell cleavage.

#### Analysis of Bouncer localization during egg activation

To analyze the localization of Bouncer in zebrafish eggs during activation, un-activated, mature eggs were collected from female fish expressing sfGFP-Bouncer as well as membrane-bound lynTomato (tg[*ubi:sfGFP-bncr, cmlc2:GFP*]; tg[*actb2:lyn-Tomato*]) by squeezing. Eggs were mounted in a petri dish with cone-shaped agarose molds (1.5% agarose in E3 medium) filled with Hank’s saline (0.137 M NaCl, 5.4 mM KCl, 0.25 mM Na_2_HPO_4_, 1.3 mM CaCl_2_, 1.0 mM MgSO_4_, 4.2 mM NaHCO_3_) to slow down activation. Imaging was performed at a LSM700 Axio Imager upright system (Zeiss) with a 40×0.75 Archoplan dipping lens at room temperature. FIJI was used to measure the fluorescence intensity of sfGFP-Bouncer and membrane-bound lynTomato at the membrane from a distance of up to 130 μm from the micropyle.

#### Imaging of sperm approach

Sperm was squeezed from 2-4 wild-type male fish and kept in 150 μl Hank’s saline containing 0.5 μM MitoTracker™ Deep Red FM (Molecular Probes) for >10 minutes on ice. Un-activated, mature eggs were obtained by squeezing a female wild-type or *bncr*^−/−^ fish. To prevent activation, eggs were kept in sorting medium (Leibovitz’s medium, 0.5 % BSA, pH 9) (*52*) at room temperature. The eggs were kept in place using a petri dish with cone-shaped agarose molds (1.5% agarose in sorting medium) filled with sorting medium. Imaging was performed at a LSM700 Axio Imager upright system (Zeiss) with a 10×/0.3 Archoplan dipping lens (781.96 msec interval, for 1-5 minutes). To start fertilization, the sorting medium was removed without letting the dipping lens go dry and at least 1 ml of E3 medium was carefully added close to the egg in order to induce activation. Immediately afterwards, 10 μl of the stained sperm was added as close to the egg as possible. The resulting timelapse movies were analyzed using FIJI.

#### Analysis of sperm-egg binding

Sperm was squeezed from 2-4 male fish and kept in 100 μl Hank’s saline + 0.5 μM MitoTracker™ Deep Red FM on ice. Un-activated, mature eggs were squeezed from a wild-type or *bncr*^−/−^ female fish and activated by addition of E3 medium. After 10 minutes, 30 wild-type or Bouncer-deficient eggs were manually dechorionated using forceps. The dechorionated eggs were transferred to a glass dish and incubated at room temperature for > 1 hour. To enable sperm binding, the glass dish was tilted and as much E3 medium as possible was removed without letting the eggs become dry, 45 μl of the sperm solution was distributed onto each egg batch and the sperm-egg mixture was incubated for another 2 minutes. The glass dish was then carefully filled with E3 medium and the eggs were transferred to a zebrafish transplantation mold (Adaptive Science Tools) (1.5% agarose in blue water) to hold them in place during imaging. Imaging was performed at a LSM700 Axio Imager upright system (Zeiss) with a 10×/0.3 Archoplan dipping lens. Sperm attached to the cell of the egg were counted during live imaging. If possible, single sperm were counted (1-10 sperm); if more than 10 sperm were attached to the egg, the egg was classified as >10 sperm bound.

#### Analysis of gDNA content in zebrafish, medaka and hybrids

Zebrafish, medaka, and hybrid gDNA was isolated according to standard protocols (*53*). In brief, a single embryo at sphere stage (zebrafish, hybrids) or stage 11-12 (medaka) was transferred into 100 μl 50 mM NaOH and incubated for 20 min at 95 °C (the medaka embryos were disrupted with a pipette tip prior to incubation). The samples were cooled to 10°C, and 1 μl of the isolated gDNA was used for a touchdown PCR (protocol: 95 °C for 2 min, followed by 15 cycles of: 95 °C for 30 sec, 70 °C for 30 sec (−1 °C each cycle), 72 °C for 40 sec, followed by 20 cycles of: 95 °C for 30 sec, 55 °C for 30 sec, 72 °C for 40 sec, followed by 72 °C for 5 min and hold at 10 °C) (*54*) using Taq polymerase and species-specific primers (zebrafish: z-vox, z-klf4; medaka: m-vox, m-klf4).

#### Statistical analysis

Statistical analysis was performed with GraphPad Prism 7 software. Statistical significance of differences between two experimental groups was assessed wherever applicable by either a two-tailed Student’s t-test if the variances were not significantly different according to the F test, or using a non-parametric test (Mann-Whitney or Kruskal-Wallis with Dunn’s test for multiple comparisons) if the variances were significantly different (p < 0.05). Differences between two data sets were considered significant at p < 0.05 (*) (p < 0.01 (**); p < 0.001 (***); n.s., not significant).

#### Data availability

RNA-Seq data first reported here were deposited at the Gene Expression Omnibus (GEO) and is available under GEO acquisition number GSE111882.

#### Primer sequences

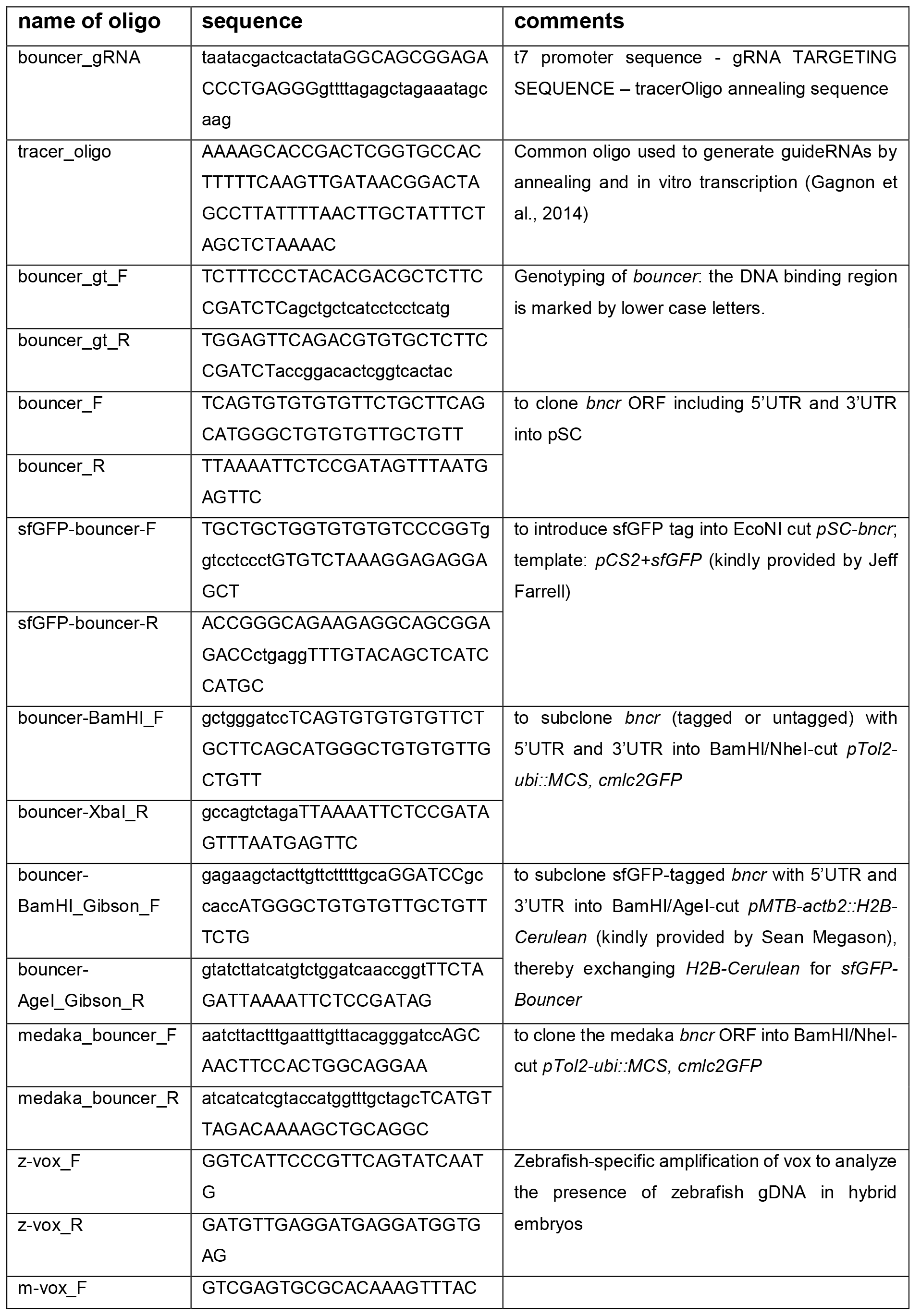

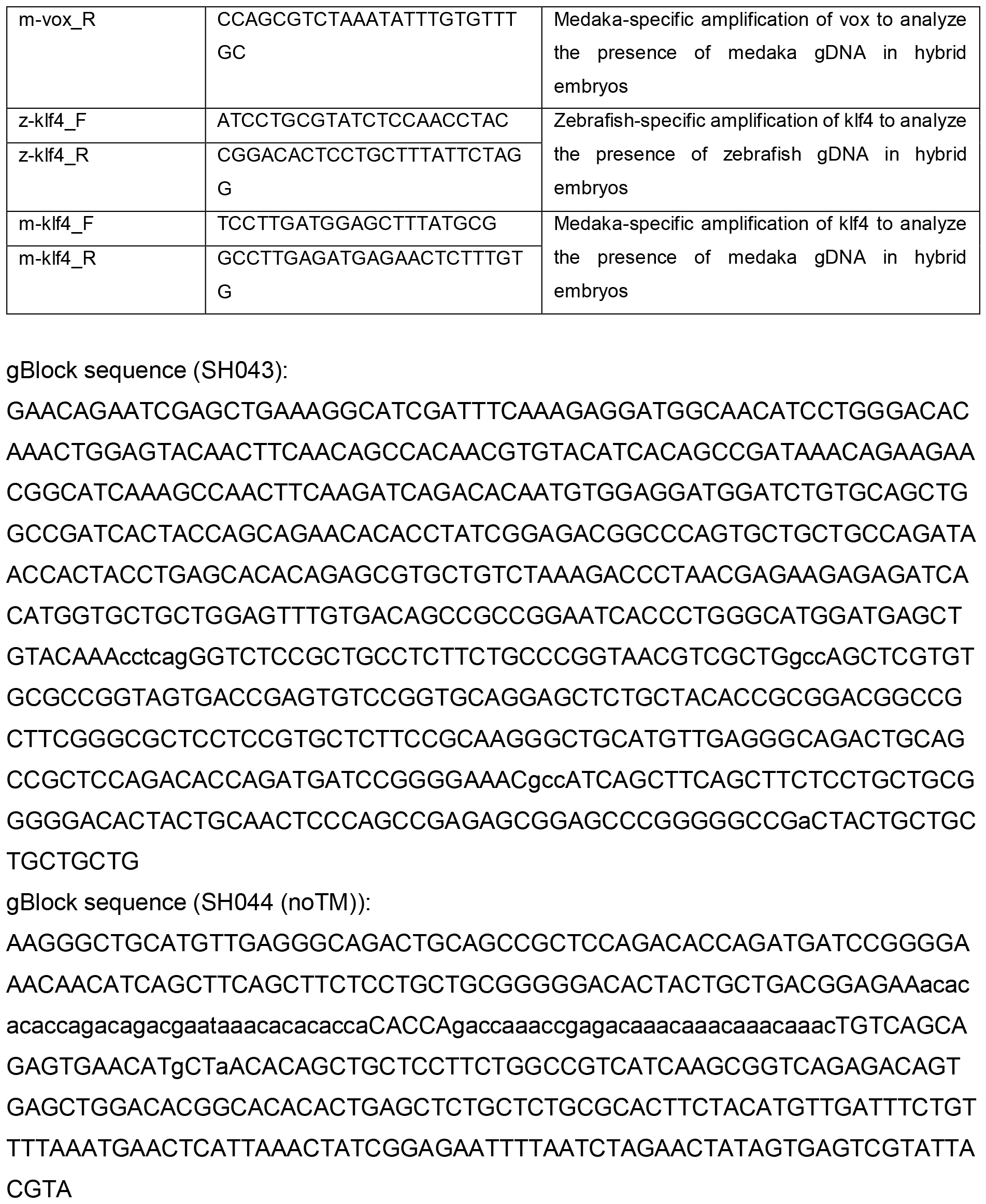

**Supplementary Figure 1:**
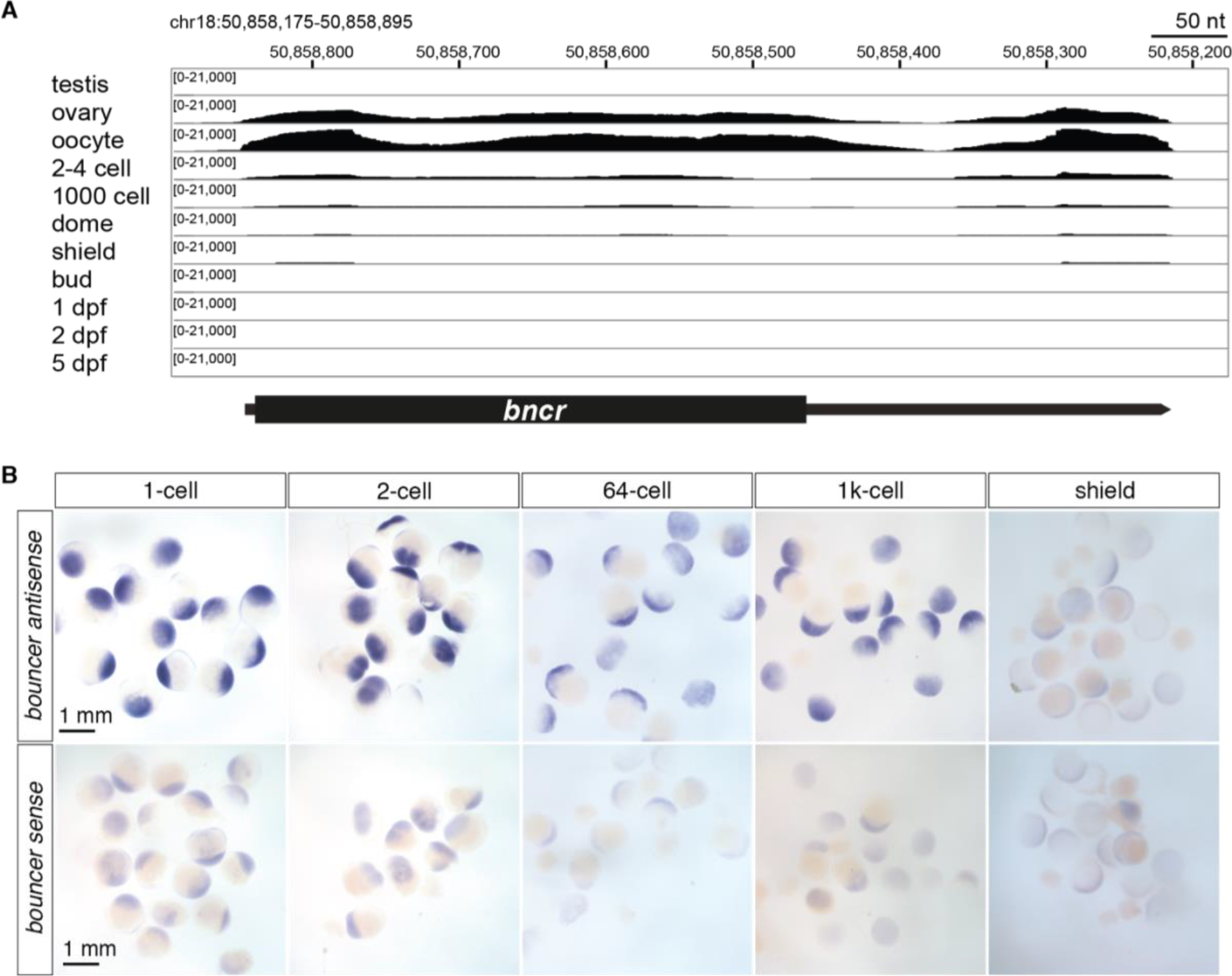
Bouncer is maternally provided. A – Expression of zebrafish *bouncer* in the germline and during embryonic development. Coverage tracks for RNA sequencing ((*55*) and this study) show high transcription of the single exon *bouncer* gene in the female germline, and decreasing amounts of *bouncer* RNA upon progression through embryogenesis. Genomic coordinates are based on GRCz10. Dpf, days post fertilization. B – *In situ* hybridization confirms that *bouncer* is a maternally provided transcript. Shown are representative overview images of *in situ* hybridization using an antisense *bouncer* probe (top) and a sense *bouncer* probe (bottom; negative control for unspecific staining).

**Supplementary Figure 2:**
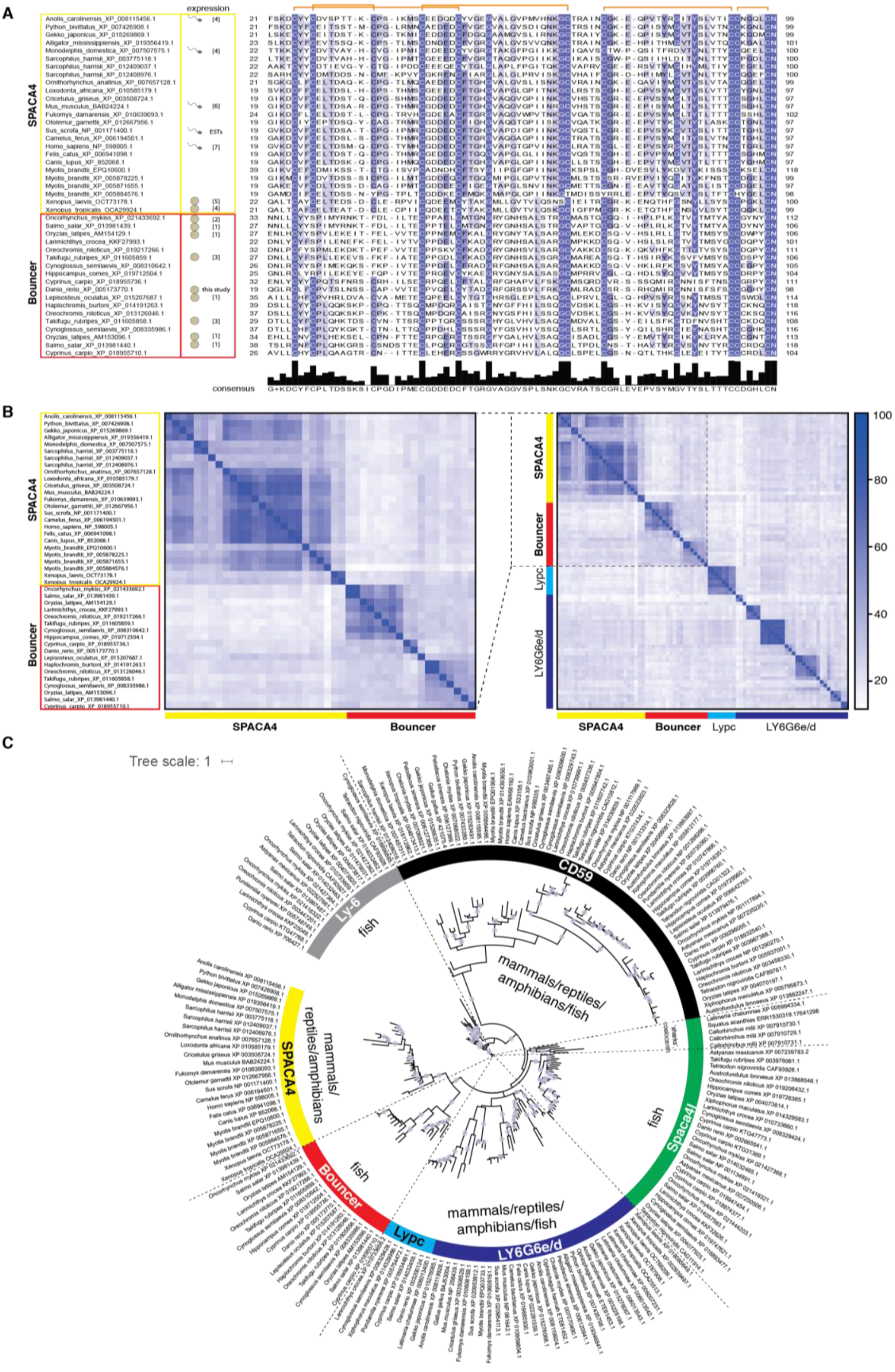
Bouncer/SPACA4 belongs to the divergent Ly6/uPAR protein family. A – Alignment of representative protein sequences of fish Bouncers and homologous mammalian and reptile SPACA4s shows high sequence divergence between species, apart from the well conserved cysteines. Only amino acid sequences of the conserved mature domain are shown; the extent of the mature domain displayed here is based on the prediction for zebrafish Bouncer. Predicted disulfide bridges are indicated in orange. Sex-specific germline expression is indicated for organisms where expression data is available (sperm: testis-specific expression; egg: ovary-specific expression). (1) (*30*) (RNA-Seq data from *Salmo trutta*); (2) (*56*); (3) (*32*); (4) (*29*); (5) (*31*); (6) (*28*); (7) human expression based on ebi-baseline expression atlas (GTEx Portal); ESTs are derived from NCBI. For sequences and accessions see Supplementary Dataset 1 and 2 and Supplementary Table. B – Bouncer/SPACA4 proteins have divergent amino acid sequences. Left: Heat map showing the percentage amino acid identity between Bouncer and SPACA4 proteins of the subset of vertebrate species shown in A. Right: Heat map showing the percentage amino acid identity between Bouncers, SPACA4, Lypc, and the group of LY6G6e/d proteins. C – Phylogenetic tree of Ly6/uPAR proteins that are most similar to the SPACA4/Bouncer family. The tree was constructed with a selected subset of vertebrate species (see Materials and Methods for details). Bootstrap values of ≥95 are indicated by filled purple circles. The color-coded group classifications (SPACA4, Bouncer, Lypc, LY6G6e/d, Spaca4l, CD59, Ly-6) follow the human, mouse or zebrafish protein nomenclature.

**Supplementary Figure 3:**
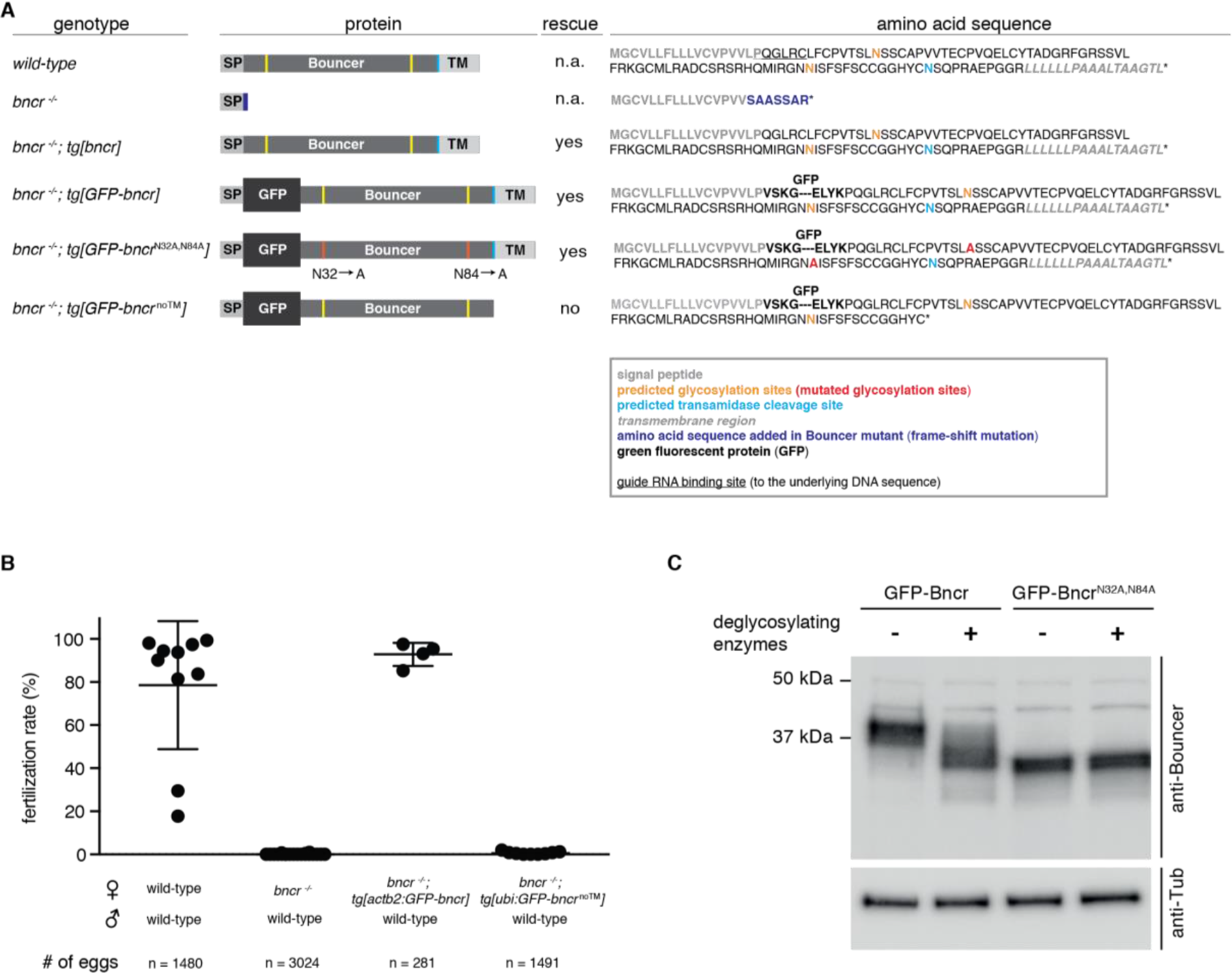
Rescue ability of Bouncer protein variants. A – Schematic representation of Bouncer protein variants and their rescuing ability of *bncr*^−/−^ females. The schemes indicate the signal peptide (SP), the two predicted N-glycosylation sites (yellow or red (mutated)), the transamidase cleavage site (turquoise) and the C-terminal transmembrane region (TM). Orange brackets indicate predicted disulfide bonds. Annotated full-length amino acid sequences are shown on the right. Note that for GFP, only the first and last four amino acids are shown. Wild-type: endogenous Bouncer; *bncr*^−/−^: loss-of-function *bouncer* mutant (13-nt deletion after the signal peptide sequence); *tg[bncr]*: transgenically expressed Bouncer; *tg[GFP-bncr]*: transgenically expressed GFP-tagged Bouncer; *tg[GFP-bncr*^*N32A,N84A*^]: transgenically expressed GFP-tagged non-glycosylatable Bouncer; *tg[GFP-bncr*^*noTM*^]: transgenically expressed GFP-tagged Bouncer lacking the GPI-anchoring site and the C-terminal transmembrane region. B – Quantification of the fertilization rates of the indicated crosses. n = total number of eggs. C – GFP-tagged Bouncer^N32A,N84A^ is not glycosylated. Treatment of GFP-Bncr^N32A,N84A^-containing samples with deglycosylating enzymes does not affect the molecular size of GFP-Bncr^N32A,N84A^, which is in contrast to treatment of GFP-Bncr-containing samples, which decreases in size. Twenty caps of dome stage embryos expressing GFP-Bncr or GFP-Bncr^N32A,N84A^ were loaded per lane. Bouncer was detected with an anti-Bouncer-specific antibody. Tubulin is shown as loading control.

**Supplementary Figure 4:**
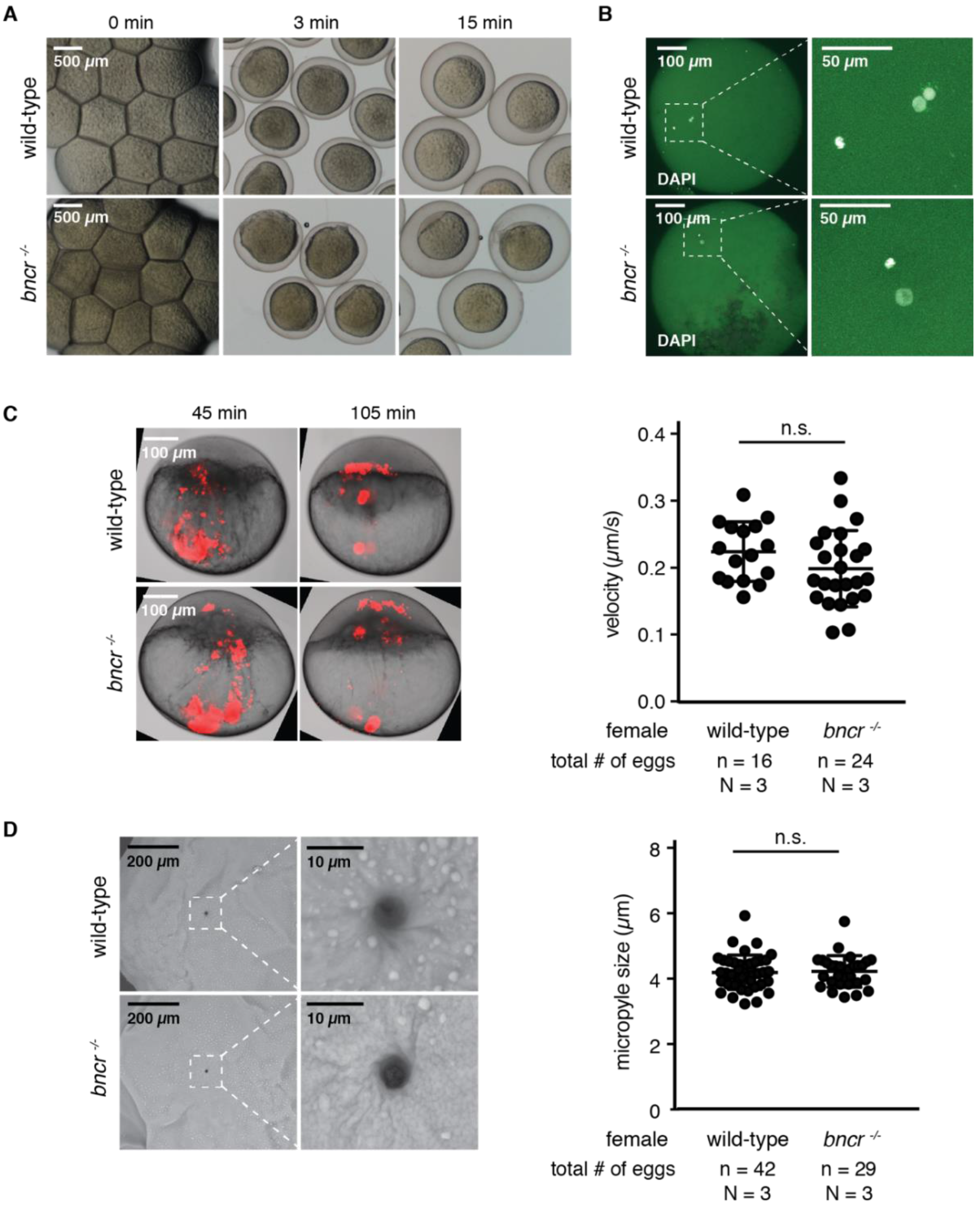
Bouncer is not required for oocyte activation. A – *Bncr*^−/−^ females produce morphologically normal eggs that elevate their chorions similar to wild-type eggs after egg activation. B – The polar body is extruded normally in Bouncer-deficient eggs. Representative image of fixed and DAPI-stained wild-type and Bouncer-depleted eggs ~15 minutes after mating. The wild-type egg contains three DAPI-stained spots (female nucleus, male nucleus, and more condensed polar body at the bottom left corner), while the Bouncer-depleted egg only contains two DAPI-stained spots (female nucleus and more condensed appearing polar body). C – Cytoplasmic streaming is normal in *bncr*^−/−^ eggs. Fluorescent beads were injected into wild-type or Bouncer-deficient eggs, and bead movement was tracked. Left: Representative images. Right: Quantification of bead velocity. Each data point represents the average velocity of all tracked beads in one egg. Mean and standard deviation are indicated (p=0.141, unpaired t-test; n = total number of eggs; N = number of biological replicates). D – The micropyle is present and of normal size in *bncr*^−/−^ eggs. Left: Representative scanning electron microscopy images of unfertilized wild-type and *bncr*^−/−^ eggs 30 seconds after activation, showing the presence of the micropyle in Bouncer-deficient eggs. Right: Quantification of the size of the micropyle based on analysis of Coomassie-stained chorions. Mean and standard deviation are indicated (p = 0.804, unpaired t-test; n = total number of eggs; N = number of biological replicates).

**Supplementary Figure 5:**
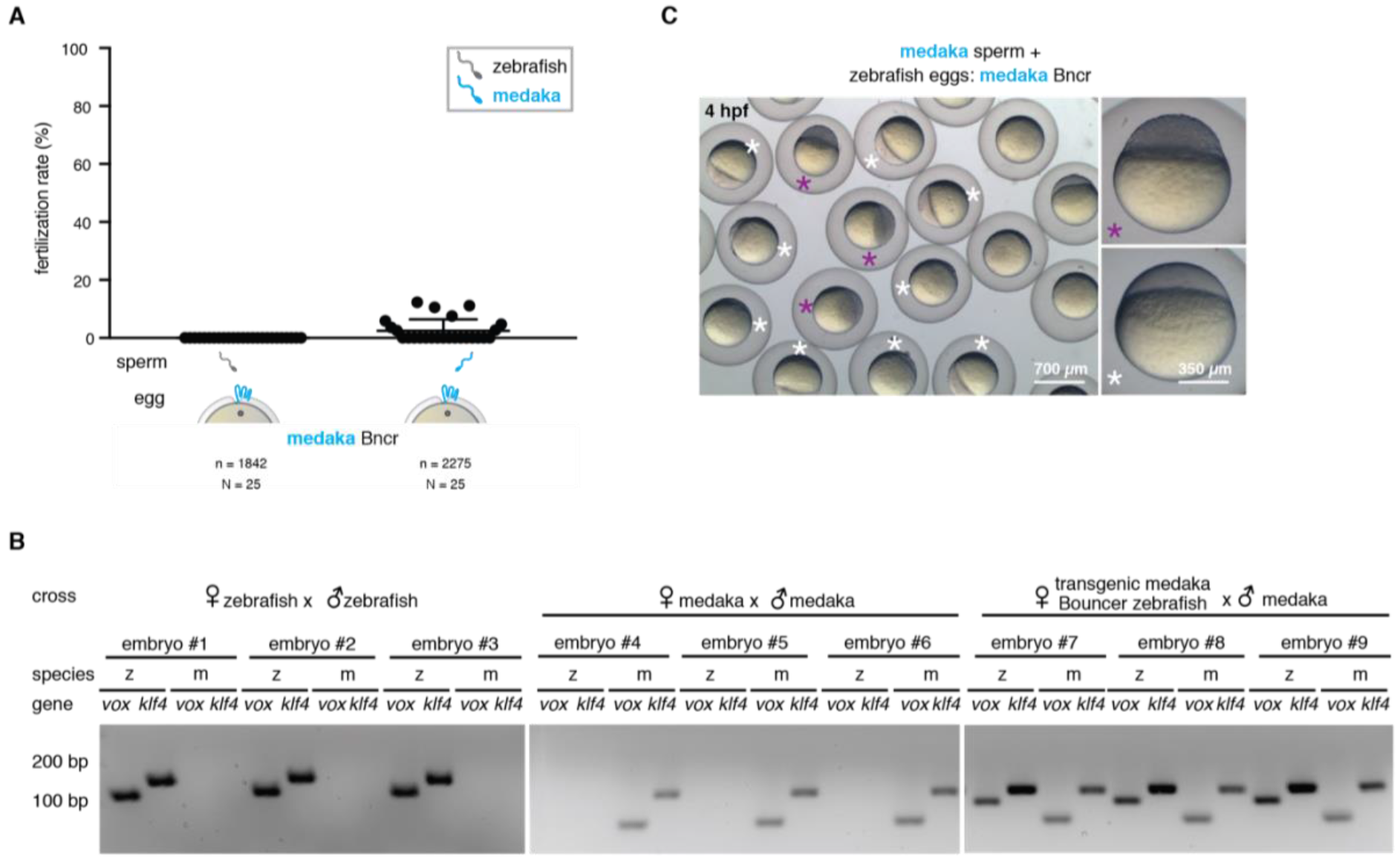
Zebrafish eggs expressing medaka Bouncer can be fertilized by medaka sperm and develop into hybrid embryos. A – Medaka Bouncer is sufficient to allow entry of medaka sperm into zebrafish eggs. While zebrafish sperm was unable to enter zebrafish *bncr*^−/−^ eggs expressing medaka Bouncer, *in vitro* fertilization with medaka sperm resulted in an average fertilization rate of 2.4%. Data is shown for all 17 medaka Bouncer-expressing females tested. Mean and standard deviation are indicated. n = total number of eggs; N = number of biological replicates. B – Fertilization of zebrafish eggs expressing only medaka Bouncer by medaka sperm results in hybrid embryos. Both zebrafish and medaka genomic DNAs (gDNAs) are present in hybrid embryos. gDNAs of single zebrafish embryos (#1-3), medaka embryos (#4-6), and hybrid embryos (#7-9) were isolated at sphere stage (zebrafish and hybrid) or between stages 11-12 (medaka) and used for PCR. Zebrafish- and medaka-specific primers (indicated as “z” or “m”) were used to amplify *vox*- and *klf4*-specific gene regions. C – Fertilization of zebrafish eggs expressing only medaka Bouncer yields medaka-zebrafish hybrid embryos that undergo gastrulation movements. All images were taken at four hours post-fertilization (hpf). White asterisks indicate unfertilized embryos (resembling one-cell stage embryos), and purple asterisks indicate fertilized embryos.

### Supplementary Movie 1

Cytoplasmic streaming is normal in Bouncer-depleted eggs. Shown are representative time-lapse series for a wild-type and *bncr*^−/−^ embryo that were injected with FluoroSpheres immediately after egg activation.

### Supplementary Movie 2

Sperm can approach the micropyle area in Bouncer-depleted eggs. Shown are representative time-lapse series for MitoTracker-labelled wild-type sperm (magenta) approaching the micropyle area of a wild-type and a *bncr*^−/−^ egg. Imaging started immediately after sperm addition.

### Supplementary Table

Table with NCBI protein and cDNA IDs, sorted according to the phylogenetic tree (192 entries). Columns contain the following information: group (SPACA4, Bouncer, Lypc, LY6G6d/e, Spaca4l, CD59, Ly-6); ID in figures (in protein sequence alignment and phylogenetic tree); species; NCBI protein accession number (if available); NCBI nucleotide (mRNA or genomic) accession number (if available); description (retrieved from the NCBI protein entry); comment on coding sequence retrieval. *Oryzias latipes* sequences were retrieved by conceptual translation, and the sequence of *Squalus acanthias* was retrieved by contig assembly and conceptual translation.

### Supplementary Dataset 1

Coding sequences of members of the Ly6/uPAR family (192 entries)

### Supplementary Dataset 2

Protein sequences of members of the Ly6/uPAR family (192 entries)

